# Actin Depolymerization Factor (ADF) Moonlighting: Nuclear Immune Regulation by Interacting with WRKY Transcription Factors and Shaping the Transcriptome

**DOI:** 10.1101/2025.04.29.651294

**Authors:** Pai Li, Brittni Kelley, Zizhang Li, Bruce Proctor, Alex Corrion, Xuan Xie, Ryan Sheick, Yi-ju Lu, Mika Nomoto, Cheng-i Wei, Yasuomi Tada, Sheng-Yang He, Shunyuan Xiao, Brad Day

**Affiliations:** Department of Plant, Soil and Microbial Sciences, Michigan State University, East Lansing, MI 48824, USA; Department of Plant Biology, Michigan State University, East Lansing, MI 48824, USA; Institute for Bioscience and Biotechnology Research, University of Maryland, Rockville, MD 20850, USA; Plant Resilience Institute, Michigan State University, MI 48824, USA; Lyman Briggs College, Michigan State University, MI 48824, USA; Cell & Molecular Biology Program, Institute for Quantitative Health Science and Engineering, Michigan State University, East Lansing, MI 48824, USA; MSU-DOE Plant Research Laboratory, Michigan State University, East Lansing, MI 48824, USA; The Department of Plant Pathology and Microbiology, National Taiwan University, Taipei, 10617, Taiwan; Center for Gene Research, Nagoya University, Furo-cho, Chikusa-ku, Nagoya, Aichi 464-8602, Japan; Department of Nutrition and Food Science, University of Maryland College Park, MD 20742; Howard Hughes Medical Institute, Duke University, Durham, NC 27708, USA; Department of Biology, Duke University, Durham, NC 27708, USA; Department of Plant Sciences and Landscape Architecture, University of Maryland College Park, MD 20742, USA; Research and Innovation, University of Tennessee–Knoxville, Knoxville, TN 37996, USA

## Abstract

Remodeling of the actin cytoskeleton is a critical process for plant immunity, essential for the transport, activation, and stabilization of immune-regulatory molecules and organelles. In this process, actin depolymerization factors (ADFs) function as key players through severing and depolymerizing actin microfilaments. However, recent evidence suggests that ADFs may possess non-canonical immune functions inside the nucleus, in addition to the canonic cytosolic role, a phenomenon not adequately explained by the traditional mechanistic model of ADF-actin dynamics. In this study, we demonstrate that Arabidopsis ADFs exhibit a moonlighting function in the nucleus, where they interact with transcriptional machinery to regulate the transcriptome during both the resting state and the immune responses. We show that ADF2/3/4 have redundant functions in defense against virulent and avirulent *Pseudomonas syringae*. Notably, it is nuclear – rather than cytosolic – ADFs that contribute to defense against *P. syringae* and mediate pro-immune transcription. Mechanistically, we demonstrate that nuclear ADFs interact with transcription factors, histone complexes, and other components of the transcriptional machinery. Specifically, ADF2/3/4 can form a complex with WRKY transcription factors, such as WRKY22/29/48, thereby directly regulating WRKY activity to shape the pro-immune transcriptome. In summary, our study reveals that ADFs moonlight as direct regulators of transcription factors, mediating a broad range of nuclear-cytoplasmic regulation in plant immunity and potentially other biological processes.

## Introduction

Plants have evolved sophisticated immune systems to survey their environment, recognize potential threats, and mount specific defensive responses. Over the past three decades, significant progress has been made in identifying and characterizing core immune signaling mechanisms, from pathogen perception and signal transduction to transcriptional reprogramming and downstream biochemical and morphological defenses (Jones et al. 2024). However, plant immunity is not a reductionist, single-thread process. An individual pathogen can introduce multiple pathogen-associated molecular patterns (PAMPs) and avirulence (Avr) effectors, activating both PAMP-triggered immunity (PTI) and effector-triggered immunity (ETI) in a dynamic, densely intertwined network across temporal and spatial scales (Yuan et al. 2021). While viewing immunity as a complex network provides a realistic perspective, it pushes a reductionist question to the foreground – *What central processes are orchestrating this entire immune network?*

Recently, a core participant of the conceptual immune signaling network has emerged in the shape of physical molecular network – the cytoskeleton – which presents a systemic and integral perspective towards a fuller understanding of plant immunity (Li and Day 2019; Wang et al. 2022; Lu et al. 2023; Sinha et al. 2024). To date, five types of cytoskeleton have been identified in eukaryotes: actin, microtubule, intermediate filament (Pollard and Goldman 2018), septin (Van Ngo and Mostowy 2019), and spectrin (Teliska and Rasband 2021), among which plants have two major types: actin and microtubules. Both are constructed by polymerized monomers that form filament meshwork that provides mechanical support to maintain cell shape and enable cellular and intracellular mobility. As one of the most important features of cytoskeleton, the architecture of cytoskeleton is highly dynamic, as both polymerization and depolymerization repetitively occur simultaneously towards an equilibrium. Therefore, this feature enables plants to actively re-model the cytoskeletal architecture to mediate desired cellular and physiological responses upon various signals, including immune activation. For example, inoculation of plants with virulent *Pseudomonas syringae* pv. tomato DC3000 (abbr. DC3000) has been shown to lead to increases in the density and bundling of actin filament of Arabidopsis epidermis cell in a global pattern (Li et al. 2022). In the case of filamentous pathogen, actin remodeling is usually observed as a condensed plate adjacent to haustorium or attempt penetration site (Qin et al. 2021; Sharma and Chandran 2022). Consequently, cytoskeletal remodeling is regarded as essential for both immunity and pathogenesis, as interfering cytoskeletal dynamics by cytoskeleton-targeted chemicals and effectors commonly leads to alternated resistance phenotype in various cases (Li and Day 2019). At the mechanism level, these phenotypes are generally attributed to the cytoskeleton’s roles in scaffolding and stabilizing plasma membrane-localized immune complexes and directing the transport of pro-immune molecules and organelles. (Li and Day 2019; Wang et al. 2022; Lu et al. 2023; Sinha et al. 2024).

Polymerization and depolymerization of actin filaments is facilitated by profilins and ADFs/cofilins, respectively (Lappalainen et al. 2022), and disabling the activity of either results in dampened immune signaling (Li and Day 2019). In the case of ADFs, multiple pieces of genetic evidence have demonstrated that specific ADFs in plants (e.g., Arabidopsis, barley, cotton, cucumber, common bean, and wheat) can regulate resistance to various types of pathogen, pests, and commensals (Sun et al. 2023). In most of these studies, while direct and detailed molecular mechanisms are lacking, it is generally accepted that the altered resistance phenotypes are attributed to the abnormal actin dynamics due to lack of ADF-driven actin depolymerization, a key step in actin remodeling in response to multiple immune signals (Li and Day 2019; Sun et al. 2023). However, a recent study from Inada et al. provides a distinctive insight into the relationship between ADF and immune signaling, which revealed that Arabidopsis subclass I ADFs (ADF1/2/3/4) contribute to the susceptibility to powdery mildew exclusively when these ADFs enter the nucleus (Inada et al. 2016). Because previous knowledge generally assumes, as noted above, that actin and its associated proteins (such as ADFs) facilitate immune process in the form of “canonic cytoskeleton” in the cytosol, the study by Inada and colleagues introduced an intriguing question related to the role of actin in plant immunity, or even general cell biology: *Does ADF, traditionally associated to cytosolic actin dynamics, perform a moonlighting molecular function within the nucleus?*

The dynamic turnover of actin filaments is regulated by a suite of actin-binding proteins, including profilins, which promote polymerization, and ADF/cofilins, which facilitate depolymerization (Lappalainen et al., 2022). Genetic studies across multiple plant species have shown that ADFs are critical for resistance against a wide range of pathogens, pests, and commensals (Sun et al., 2023). These findings are typically explained by the canonical model where ADF-driven depolymerization is a prerequisite for the actin remodeling required for effective immunity (Li and Day, 2019; Sun et al., 2023). However, this model does not account for all observations. A study by Inada et al. (2016) revealed that subclass I ADFs in Arabidopsis (ADF1/2/3/4) contribute to susceptibility to powdery mildew, a function executed exclusively within the nucleus. This finding challenges the conventional cytosol-centric view of ADF function and raises an intriguing question: Do ADFs perform a “moonlighting” function in the nucleus, independent of their canonical role in the cytoplasm?

In plants, the nuclear functions of ADF and actin are largely unknown. Conversely, studies in mammalian systems have shed light on potential mechanism(s) that engage actin and its dynamics in the nucleus. While predominantly localized in the cytosol, both mammalian actin and ADF have been shown to possess nuclear-localized roles. To enable this, cofilin, a member of ADF family, shuttles actin monomers (ACT) into the nucleus as cofilin-ACT complex, via a nuclear localization signal (NLS) recognized by importin-9 (Dopie et al. 2012). Inside the nucleus, actin also exhibits morphological dynamic processes, partially resembling cytosolic actin cytoskeleton but with unique features. For example, nuclear actin remodel into nod-like structures upon chemical/environmental stress such hypoxia, pathogens, and drugs (Kloc et al. 2021), which is mechanistically associated with actin’s interaction with RNA polymerases and chromatin remodeling complexes, a demonstrated role in transcriptional regulation (Kyheröinen and Vartiainen 2020; Wei et al. 2020). Interestingly, while no evidence yet suggests that canonic ADF can directly interact with transcriptional machinery without actin as the scaffold, it is reported that human Drebrin – an actin binding protein – utilizes its N-terminal ADF-H domain to interact with chromatin reader ZMYND8 to regulate its function (Yao et al. 2017). Finally, in the context of the actin-immunity relationship, it is noteworthy that ADF phosphorylation may play a role in its proposed nuclear function. The mammalian cofilin is tightly regulated by phosphorylation (Xing et al. 2024), similar to ADFs of plants (Lu et al. 2020), whose phosphorylation, denoting a pro-immune phase, generally inhibits ADF-ACT interaction. In this manner, phosphorylation can potentially release ADF from actin to promote a different type of molecular function in the nucleus. Collectively, the evidence above supports the hypothesis that ADFs possess an as-yet-undiscovered transcriptional regulatory function that contributes to plant immunity.

In this study, we investigated the biological significance of nuclear ADFs and identified the historically enigmatic “ADF moonlighting function” as transcription factor (TF) regulators in plant immunity. In brief, Arabidopsis ADF2/3/4 contribute to defense through direct interaction with certain members of the WRKY and potentially other components of the transcriptional machinery in the nucleus. Specifically, ADF4 associates with WRKY29/48, forming an ADF-WRKY-DNA complex on the promoters of defense genes, which further enhances WRKY’s activity. As a result, nuclear – but not cytosolic – ADF4 majorly contributes to defense against *Pseudomonas syringae* by facilitating transcription of genes and biological pathways related to immunity, hormone signaling, cell wall metabolism, DNA organization/replication, and other cellular processes. Besides, the role of cytosolic and nuclear ADFs in various types of plant-microbes appears to be different. Overall, we propose that the ADFs possess a moonlighting nuclear function as direct transcriptional regulators, and that both their cytosolic function (for actin dynamics) and nuclear function (for transcription) are engaged for orchestrating plant immunity and potentially other biological processes.

## Results

### Class I ADFs are expressed in leaves and redundantly contribute to defense

Our previous study reported that an *adf4* mutant displays enhanced susceptibility to *Pseudomonas syringae* pv. *tomato* (*Pst*) DC3000 harboring the avirulent effector gene *AvrPphB* (Tian et al. 2009; Porter et al. 2012). The Arabidopsis genome contains 11 ADF-encoding genes. Because all the ADF family members are small single-domain proteins without major structural differences, we hypothesized that certain levels of functional redundancy may conceal potential immune functions of any single *ADF* gene in conventional genetic analysis. To begin to address the potential redundant roles *ADFs* might play in the activation and signaling of plant immunity, we analyzed the expression profile of all *ADFs* during PTI and ETI using published mRNA-seq datasets [GSE85932, (Birkenbihl et al. 2017); GSE151885, (Saile et al. 2020)]. As shown in Fig. 1A-B, only *ADF1/2/3/4* (class I), *ADF5* (class III), and *ADF6* (class IV) were found to be constitutively expressed in rosette leaves, with minor induction following immune elicitation in some of the cases. Considering the close phylogenic relationship among ADF1/2/3/4 (Fig. 1A) and the results of previous studies suggesting altered immune phenotypes of *ADF1/2/3/4*-silenced lines (Porter et al. 2012; Inada et al. 2016), we inferred that construction of a high-order null mutant for class I *ADFs* would be highly effective in circumventing their potential functional redundancy and thereby revealing the contribution of these ADFs to plant immunity.

**Figure 1:**
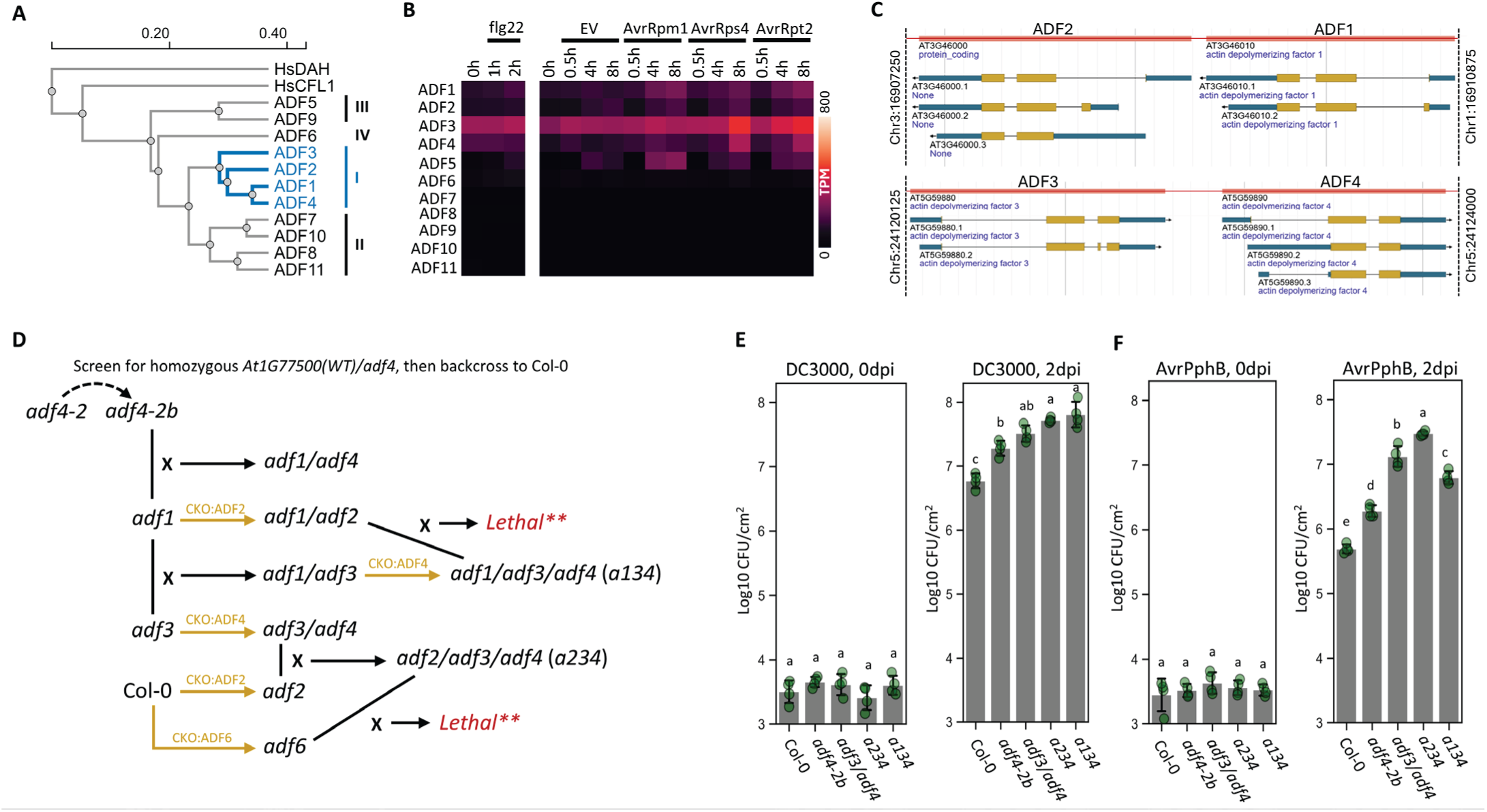
Class I ADF redundantly contributes to Resistance against Pseudomonas syringae. **A**, Phylogenic tree of Arabidopsis ADFs based on proteins sequence. Class I-IV are illustrated. Human CFL1 and Drebrin ADF-H domain (DAH) are also included, which are shown distinct from Arabidopsis ADFs. **B**, temporal expression profile of Arabidopsis ADFs in PTI and ETI. Public mRNA-seq datasets GSE85932 (PTI) and GSE151885 (ETI) were downloaded, processed, and analyzed. Expression levels are represented by TPM. **C**, genome map of ADF1/2/3/4. ADF1/2 and ADF3/4 are adjacent genes, which do not support the construction of their higher order mutants by crossing. **D**, technical routine to construct ADF high order mutants combining CRISPR and crossing. **, presumably lethal as no homozygous progenies could be identified through multiple attempts. **E**-**F**, *P. syringae* infection assay using ADF high order mutants. DC3000 (**E**) or DC3000/AvrPphB (**F**) at OD_600_ = 0.002 were infiltrated to 5-week-old Arabidopsis. Infected leaf samples were measured at 2dpi. ADF high order mutants significantly enhance the susceptibility to DC3000 and AvrPphB. Data pass ANOVA with P<0.05; non-overlapping alphabets suggest P<0.05 upon post-hoc T-tests adjusted by Benjamini–Hochberg procedure for family-wise error.

Because *ADF1/2* and *ADF3/4* are adjacent genes (Fig. 1C), constructing high order mutants by crossing is almost impossible. Therefore, we used the CRISRP-Cas9 system to introduce *adf2* and *adf4* null mutations to the *adf1* and *adf3* backgrounds respectively, which, combined with crossing, allowed us to obtain an *adf1/adf3/adf4* mutant (*a134*), and an *adf2/adf3/adf4* (*a234*) mutant (Fig. 1D; Supplemental Fig. 1). However, the quadruple mutant *adf1/adf2/adf3/adf4* was not identifiable through repetitive efforts, potentially because losing all class I ADFs is lethal or results in infertility. We also knocked out *ADF6* through CRISPR-targeted mutagenesis; similarly, *adf2/adf3/adf4/adf6* was not identifiable.

Next, to determine the genetic contribution of leaf-expressed ADFs to immunity, we challenged the high-order mutants with the virulent wild-type strain Pst DC3000 or the avirulent *Pst* DC3000 carrying *AvrPphB* to evaluate the bacterial growth (see Fig. 1E-F). In the case of DC3000, the *a134* and *a234* triple mutants showed ∼10-fold higher bacterial growth compared to Col-0, while the *adf4-2b* single mutant had only 2∼3 folds, implying that these ADFs serve overlapping or independent roles in basal immunity. When challenged with the *Pst* DC3000-*AvrPphB* strain, a similar trend was observed with the difference between *a234* and Col-0 with a greater contrast of ∼50-fold. These results suggest that Class I ADFs play positive and likely redundant and/or overlapping roles in basal defense (including PTI) and ETI against DC3000/AvrPphB.

### Nuclear ADF4 interacts with nuclear proteins including transcription machinery

As mentioned above, previous studies implied that nuclear ADF4 may play an independent role in immunity beyond its canonic function as an actin depolymerization catalyst within the cytosol (Inada et al. 2016). To explain those phenotypes, we hypothesize that ADFs can regulate the function of the transcription machinery through an unknown mechanism. To determine the potential roles of ADFs in the nucleus, we conducted a TurboID proteomics-based screening to identify potential nuclear interactors of ADF4 (Fig. 2). TurboID is engineered biotin ligase that generates unstable ATP-biotin intermediate in the presence of biotin, which immediately transfer the biotin to adjacent proteins as an affinity marker of potential interactors (Mair et al. 2019; Fig. 2B). Specifically, we adapted a engineered variant of TurboID with smaller size, namely miniTurbo (Branon et al. 2018), to create an ADF4-miniTurbo-mVenus-NLS (abbr. A4mTBYN; NLS, SV40 <u>n</u>uclear localization <u>s</u>ignal peptide) fusion protein construct, which was transformed into the *a234* background to mark potential ADF4 interactors. A fusion construct without ADF4 (abbr. mTBYN) was used as negative control (Fig. 2B). Confocal microscopy confirmed that both fusion proteins specifically localized in the nucleus, suggesting the NLS is sufficiently active (Fig. 2C, D).

**Figure 2:**
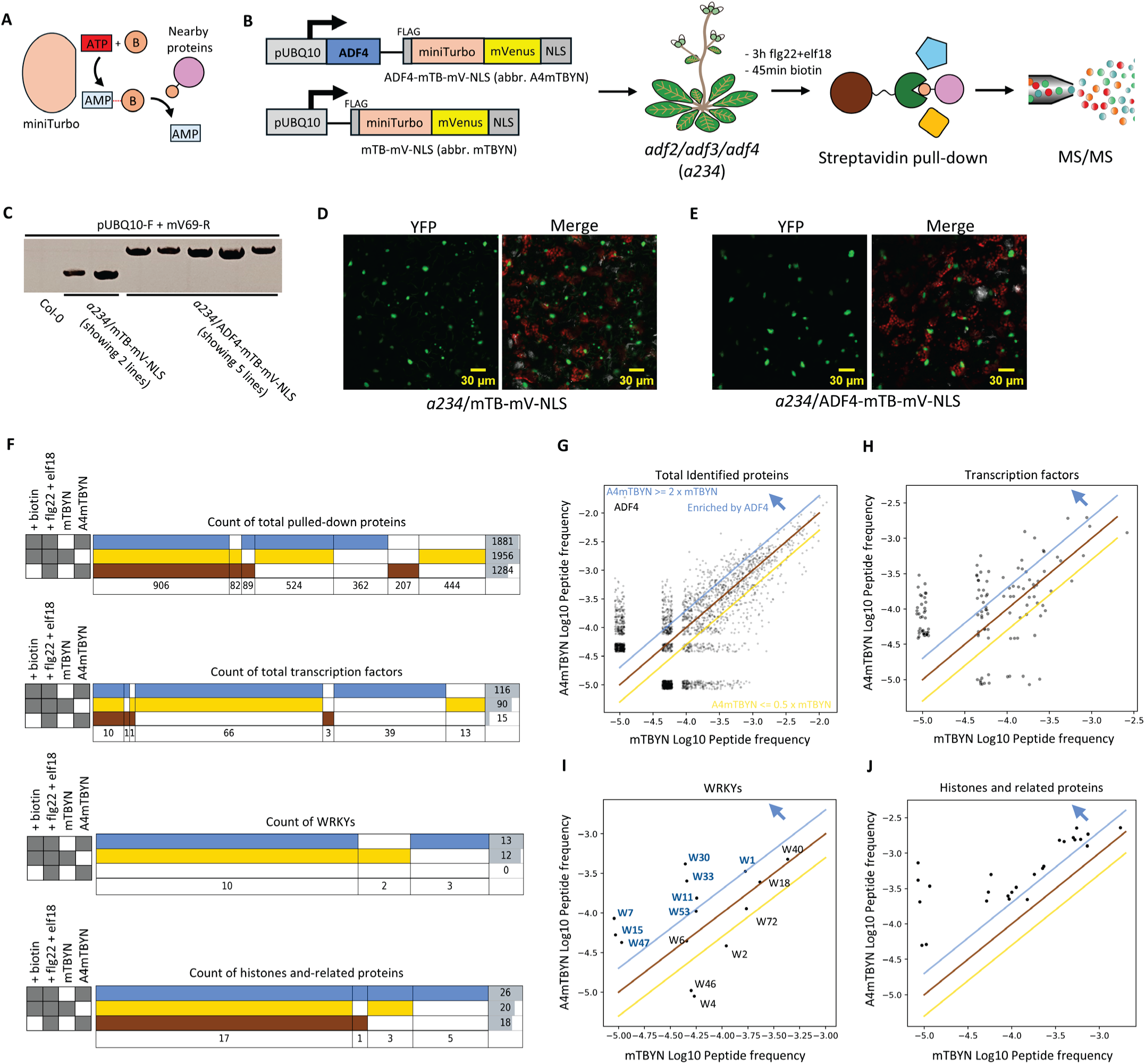
Identification of ADF4 interactors in nucleus using TurboID. **A**, TurboID mechanism. miniTurbo-fused interested protein transfers a covalent biotin tag (shown as “B”) to its nearby/interacting protein, which can be used for interactome enrichment. **B**. Experiment scheme. ADF4 was fused by miniTurbo-mVenus-NLS (mTBYN) using a flexible linker, driven by pUBQ10. This ORF was transformed into *a234* background. Plants were treated with PAMPs to induce pro-immune proteome, followed by biotin to activate labelling. ADF4-interactive in the nucleus were IP-ed and identified through MS/MS. Plants with miniTurbo-mVenus-NLS construct without ADF4 was used as the negative control. **C**, genotypic verification of TurboID constructs. **D**-**E**, subcellular localization of mTBYN (**D**) and ADF4-mTBYN (A4mTBYN; **E**) in transgenic Arabidopsis. Both exhibit exclusive nuclear localization. **F**, SuperVenn diagrams of ADF4-specific and non-specific TurboID-labelled interactome, in the order of total pull-down, transcription factors, WRKYs, and histone related proteins, from top to bottom. Note that SuperVenn is modified format of Venn diagram with flattened layout, with unlimited set number for accurate presentation. **G**-**J**, quantitative distribution of ADF4-enriched proteins. Scatter plots are used to show the relationship of logarithmic peptide frequency in mTBYN vs A4mTBYN of identified proteins, in the order of total protein (**G**), TFs (**H**), WRKYs (**I**), and Histone related proteins (**J**). Frequency of “0” is re-valued to 1e-5 to enable visualization. A point jitter is applied to visualize overlapping points. Blue line, A4mTBYN = 2 * mTBYN; brown line, A4mTBYN = mTBYN; yellow line, A4mTBYN = 0.5 * mTBYN. The distance from a top-left point to the brown line represents the protein’s enrichment level by ADF4. We define the proteins at the top-left of the blue line as the potential ADF4 interactive proteins.

In order to identify ADF4 nuclear interactors associated with immunity, we challenged the 4-5-week-old Arabidopsis rosette leaves with a mixture of flg22 and elf18 (PAMPs that elicit PTI) 3 hours before biotin treatment for 45 min, followed by streptavidin pull-down for biotin-labelled proteins. After MS/MS analysis, we identified 768 proteins with at least a 2-fold enrichment by A4mTBYN compared with mTBYN, including 362 proteins exclusively labelled by A4mTBYN (Fig. 2F, G). To zoom-in on the candidates associated with the transcriptional machinery, we investigated the candidates pool and identified 67 TFs (Tian et al. 2020) enriched by A4mTBYN, including 39 exclusive candidates (Fig 2F, H). Among those TFs, members of the WRKY family were of immediate interest, as WRKYs are generally related to stress response (including immune events) exclusively existing in plant species (Javed and Gao 2023). There was a total of 8 WRKYs specifically enriched by A4mTBYN (Fig 2F, I). Furthermore, inspired by the finding that HsDrebrin interacts with a histone reader HsZMYND8 via its ADF-H domain (Yao et al., 2017), we also searched for histone-related proteins and identified 23 candidates enriched by A4mTBYN (Fig. 2J). These results support our hypothesis that nuclear ADFs can interact with transcriptional machinery, including WRKY TFs and histone complexes, to co-regulate gene expression during an immune response. A full list of ADF4-enriched proteins is provided as Supplemental Material 1.

### ADF2/3/4 specifically interact with WRKYs

Given the importance of WRKYs in plant immunity (Javed and Gao 2023), we hypothesized that the ADF4-WRKY interaction revealed by TurboID proteomics may represent only a portion of the ADF-WRKY interaction network in Arabidopsis. This is because Arabidopsis has 11 ADFs (Fig. 1A) and 74 WRKYs (Li et al. 2020), plus 12 ACTs (McDowell et al. 1996) possibly involved into the complex due to the nature of ADF. In order to further clarify the ADF-WRKY interaction network, we developed strategies to select appropriate representative ADF and WRKY candidates to gain further insights into this interaction network.

First, we analyzed the aforementioned mRNA-seq datasets of PTI and ETI transcriptome (Fig. 1B) to determine the expression profile of all WRKYs (Supplemental Fig. 2-3). We found that all WRKYs showed low expression levels in leaves at the naive, resting state (i.e., without immune elicitation); however, during immune signaling, dozens of WRKYs are boosted, suggesting their involvement in immunity and potential relationship with ADFs participated in this process. Next, we used two criteria to further narrow down the WRKY candidates: (1) selected WRKY candidates should cover all phylogenic clades of the entire WRKY family and (2) WRKYs reported to be associated with immunity and/or pulled down by TurboID (Fig. 2I) should be prioritized. With these criteria, we finalized 16 WRKY candidates covering 8 phylogenic groups, as summarized in Fig. 3A.

**Figure 3:**
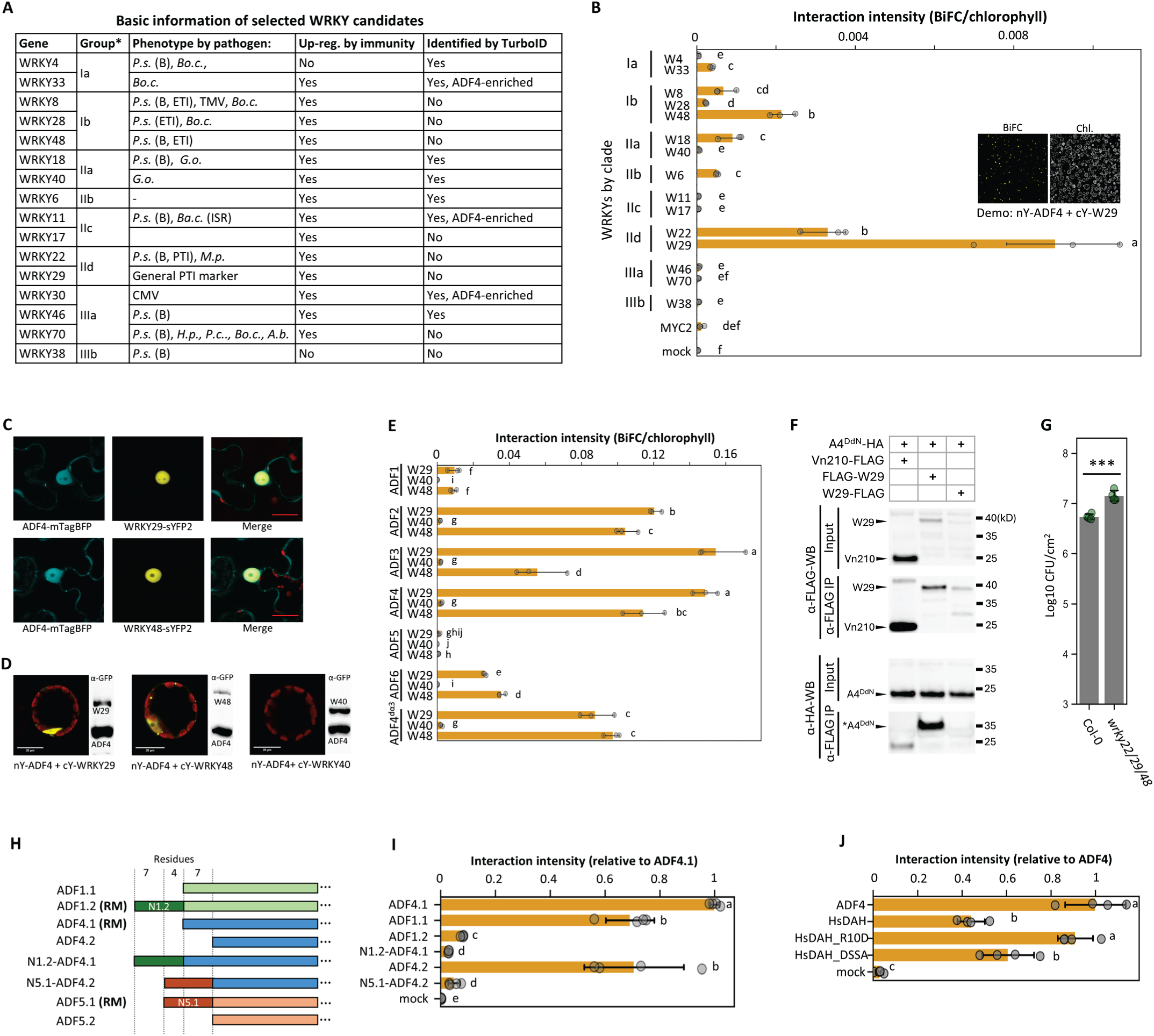
ADFs interact with WRKYs specifically. **A**, information of selected WRKY candidates. Arabidopsis WRKYs were selected to test their capability to interact with ADFs, based on their phylogenic group, immune phenotypes, induction profile, and TurboID identification. *A.b.*, *Alternaria brassicicola*; *Ba.c*., *Bacillus cereus*; *Bo.c.*, *botrytis cinerea*; *G.o.*, *Golovinomyces orontii*; *M.p.*, *Myzus persicae*; *P.c.*, *Pectobacterium carotovorum*; *P.s.*, *P. syringae*; CMV, cucumber mosaic virus; TMV, tobacco mosaic virus; (B), basal defense; (ISR), Induced systemic resistance. **B**, ADF4-interaction intensity of selected WRKYs by semi-quantitative BiFC. Controlled amount of nY-ADF4 and cY-WRKYs were transformed into Col-0 protoplast. After 12h, large scale confocal images, shown as demo, were taken for computational analysis. WRKY22/29/48 show strong interaction with ADF4. **C**, subcellular localization of ADF4 and WRKY29 in *N. benthiamiana*. **D**, single cell image of ADF4-WRKYs BiFC. Most interaction occurs in the nucleus with minor signals in vesicle-like structures. Respective immunoblots confirmed that similar amounts of ADF and WRKYs were produced for the experiment system. **E**, Arabidopsis ADFs have varied interaction intensity to WRKYs. For BiFC, controlled amount of nY-ADFs were co-transformed with cY-WRKY29/48, or cY-WRKY40 as a negative control. ADF2/3/4 have strong interaction with WRKY29/48. Actin-non-interactive ADF4 mutant, ADF4^dα3^, still maintains WRKY interaction activity. **F**, Co-IP demonstrating ADF4-WRKY29 interaction. ADF4^S6D/dα3^-NLS (abbr. A4^DdN^) was used as an alternative of wild type ADF4, to eliminate actin binding competition and improve WRKY affinity. mVenus1-210 (abbr. Vn210), the n-terminal nuclear permeable fraction of mVenus, was used as a negative control. FLAG-WRKY29 pulled-down an unknown modified version of A4^DdN^-HA (marked as “*”), but WRKY29-FLAG cannot mediate any interaction. Extremely strong expression of Vn210 introduced a weak non-specific interaction with A4^DdN^. **G**, *P. syringae* DC3000 infection assay on triple mutant *wrky22/29/48. wrky22/29/48* has dampened resistance, phenocopying *a234* (Fig. 1E). **H**, Alternative splicing of ADF1/4/5 models on the N-terminus and design of their chimera. ADF1/4/5 alternative splicing models from TAIR are aligned according to protein sequence homogeneity. The first 11 AA of ADF1.2 are fused to ADF4.1, making N1.2-ADF4.1N; the first 11 AA of ADF5.1 are fused to ADF4.2, making N5.1-ADF4.2. RM, TAIR representative gene model. **I**, N-terminal sequence of ADFs significantly impact ADF-WRKY interaction. Different transcription models of ADFs and their chimeras were co-transformed with cY-WRKY29 for BiFC. N1.2 inhibits the interaction of ADF1.1/4.1 and WRKKY29; N5.1 inhibits the interaction of ADF4.2. **J**, human drebrin ADF domain (HsDAH) interacts with WRKY29 using an interface distinguished from that of HsDAH-ZMYND8 interaction. HsDAH, and its single mutant R10D and quadruple mutant R10D/L14S/C96S/E107A (DSSA), on critical residues mediating HsDAH-ZMYND8 interface, were co-transformed with WRKY29 for BiFC. HsDAH interacts with WRKY29 with a minorly reduced intensity compared with ADF4, but R10D or DSSA mutant do not impact the interaction. All data pass ANOVA with P<0.05; non-overlapping alphabets suggest P<0.05 upon post-hoc T-tests adjusted by Benjamini–Hochberg procedure for family-wise error.

To determine which among these WRKYs mediate specific interaction with ADF, we conducted semi-quantitative BiFC to evaluate the interaction intensity of each WRKY candidate with ADF4. nY-ADF4 and cY-WRKY were co-transformed into Col-0 protoplasts to measure their overall fluorescence level using image computational approaches. As shown in Fig. 3B, half of the WRKY candidates showed detectable interaction intensity (descending group greater than “e”) with ADF4, while their interaction intensities had contrast variation. Among the candidates, WRKY22/29/48 were the strongest interactors, and WRKY6/8/18/28/33 were medium-weak interactors. Interestingly, these interactors concentrated in group Ib and IId, indicating the specificity of ADF-WRKY interaction may be related to structural features of these clades. In a further localization assay, we verified that ADF4 co-localized with WRKY29/48 in the nucleus (Fig. 3C), and their BiFC signal majorly distributed in the nucleus (Fig. 3D) as well, in accordance with their subcellular localization. Furthermore, a DC3000 infection assay using *wrky22/29/48* triple mutant showed that these WRKYs are collectively required for plant immunity (Fig. 3G), which phenocopies *a234*. Hence, we choose WRKY29 as the primary representatives for further study. Some experiments were also performed with WRKY48.

Next, to understand the interaction specificity among all leaf-expressed ADFs, we conducted BiFC on ADF1-6 in combination with WRKY29/48 as well as WRKY40 as negative control. As shown in Fig. 3E, ADF2/3/4 showed strong interaction with WRKY29/48 and no interaction with WRKY40. In comparison, ADF6 was a medium interactor; ADF1 was a weak interactor; ADF5 was a non-interactor. Since ADF1/2/3/4/6 displayed a similar interaction pattern with WRKY29/48, it is deducible that WRKYs’ interaction with these ADFs may rely on a similar mechanism. Therefore, we choose ADF4 as a representative in the following studies.

ADFs can interact with both ACT monomers and F-actin (Jaswandkar et al. 2022). To understand whether the ADF-WRKY interaction depends on ADF-ACT interaction (i.e., forming a WRKY-ADF-ACT triplex), we introduced a novel ADF4 mutant, ADF4^R98A/K100A^, referring to previous studies (Du et al. 2016; Tanaka et al. 2018). Deduced from 3D structure, this mutant is unable to interact with ACT because the two mutated residues disrupted an alpha-helix at the ACT-ADF interaction interface. We named this mutant ADF4^dα3^ (for “disrupted alpha-helix III”) and found that ADF4^dα3^ cannot bind to actin filament like general ADFs (supplemental Fig. 4). Surprisingly, ADF4^dα3^ can still strongly interact with WRKYs, indicating the ADF-WRKY interaction *per se* does not necessarily require ACT as a scaffold. Previous studies reported that immune-triggered ADF phosphorylation on ADF-ACT interface prohibits this interaction to mediate actin remodeling (Dong and Hong 2013; Lu et al. 2020). As ADF-WRKY and ADF-ACT interactions potentially use different interfaces, the phosphorylation may impact to ADF-WRKY interactions differently. To test this hypothesis, we conducted a BiFC assay using several ADF phosphomimic mutants and learned that these phosphorylation events do not substantially inhibit ADF4-WRKY29 interaction (supplemental Fig. 5). On the contrary, S6D and S105D mutations, which dissociates ADF4 from actin (Lu et al. 2020), generate slightly higher interaction intensity with WRKY29. We further confirmed the ADF4-WRKY29 interaction using co-immunoprecipitation (co-IP) in *Nicotiana benthamiana*. While wild type WRKY29 cannot pull-down significant amount of wild type ADF4 (data not shown) potentially due to competition from massive native NbACTs, we applied a modified ADF4 construct, ADF4^S6D/dα3^-NLS (abbr. A4^DdN^), to enhance ADF-WRKY interaction and eliminate ACT competition and demonstrated that A4^DdN^ indeed interacts with WRKY29 *in vitro* (Fig. 3F). These results implied that the ADF-WRKY and ADF-actin interactions are mediated by distinct interfaces and that phosphorylation, by releasing ADF from actin, may shift the equilibrium toward nuclear, WRKY-associated functions.

### N-terminal sequence of ADF regulates ADF-WRKY interaction

It was surprising to observe that ADF2/3/4, but not ADF1, displayed a strong WRKY interaction. This is because ADF1 has the closest phylogenic relationship with ADF4 (Fig. 1A, 3E). By comparing their sequences, we realized that the representative gene model ADF1.2, used in our initial experiments, has an additional 11 residue overhang on the N-terminus (namely N1.2) compared to the rest of class I ADFs, but they are ignored by the phylogenic analysis algorithm that mostly focuses on homologous regions. Therefore, the low BiFC intensity of ADF1(.2) might suggest a potential inhibitory function of ADF N-terminal sequence. To test this hypothesis, we measured WRKY interactions of different splicing products of ADF1/4 as well as their chimeras. As shown in Fig. 3H, I, ADF1.1 (the isoform without N1.2), ADF4.1, and ADF4.2 have strong interactions with WRKY29, while ADF1.2 interaction with ADF4.1 was low as expected. Further, when N1.2 was fused to ADF4.1, it is no longer able to interact with WRKY29 strongly, suggesting that N1.2 inhibits the ADF-WRKY interaction. An alternative explanation is that N1.2 is structurally rigid and increases the distance between nY and cY to bind each other. To inspect this possibility, we tested N5.1-ADF4.2 chimera using N-terminal sequence of ADF5.1, which is only 4 residues longer than ADF4.1 (Fig. 3H). As shown in Fig. 3I, N5.1-ADF4.2 chimera displayed a low WRKY interaction, indicating that it is likely the sequence and structure, rather than the length of the ADF N-terminal overhang, that regulates the ADF-WRKY interaction. It is deducible that the alternative splicing of ADFs may be functionally related to the equilibrium of their cytosolic-nuclear activity.

### ADF-WRKY and HsDrebrin-ZMYND8 have different interaction interfaces

Since *Hs*Drebrin can use its ADF-H domain to bind the histone reader ZMYND8 as a potential transcription regulatory mechanism, we were curious whether the ADF-WRKY interaction in Arabidopsis also utilizes the same interface. To answer this question, we expressed the truncated *Hs*<u>D</u>rebrin <u>A</u>DF-<u>H</u> domain (*Hs*DAH) in our BiFC system and found that *Hs*DAH had a lower, yet still robust, interaction with WRKY29 (Fig. 3J). When we introduced single or quadruple mutations of *Hs*DAH on the HsDrebrin-ZMYND8 interface (Yao et al. 2017), the interaction intensity was not substantially changed (Fig. 3J). These results indicated that ADF-WRKY and HsDrebrin-ZMYND8 interactions likely do not share similar interfaces.

### ADFs interact with WRKY-DNA complex

WRKYs regulate transcription through binding to the W-box motif ((T)TGAC(C/T)) in the promoter sequences of stress-responsive genes (Jiang et al., 2017a). Considering the ADF-WRKY interaction, we hypothesized that ADF is involved in transcriptional regulation as a component of an ADF-WRKY-DNA complex. To test this hypothesis, we conducted a chromatin immunoprecipitation (ChIP) experiment using *adf4/wrky29* double mutant protoplast co-expressing ADF4-HA and/or WRKY29-MYC (Fig. 4A). WRKYs can bind to the W boxes in their own promoters, as shown for WRKY18/40/60 (Liu et al., 2012). We found 5 W-boxes (see Fig. 4B) in the WRKY29 promoter (pW29; -1900 to +1). Additionally, a previous study showed that WRKY29 can bind the W-boxes of pBAG7, an ER-nuclear co-chaperone involved in unfolded protein response (Li et al., 2017b). Therefore, we performed ChIP assay with the W-boxes in pW29 and pBAG7.

**Figure 4:**
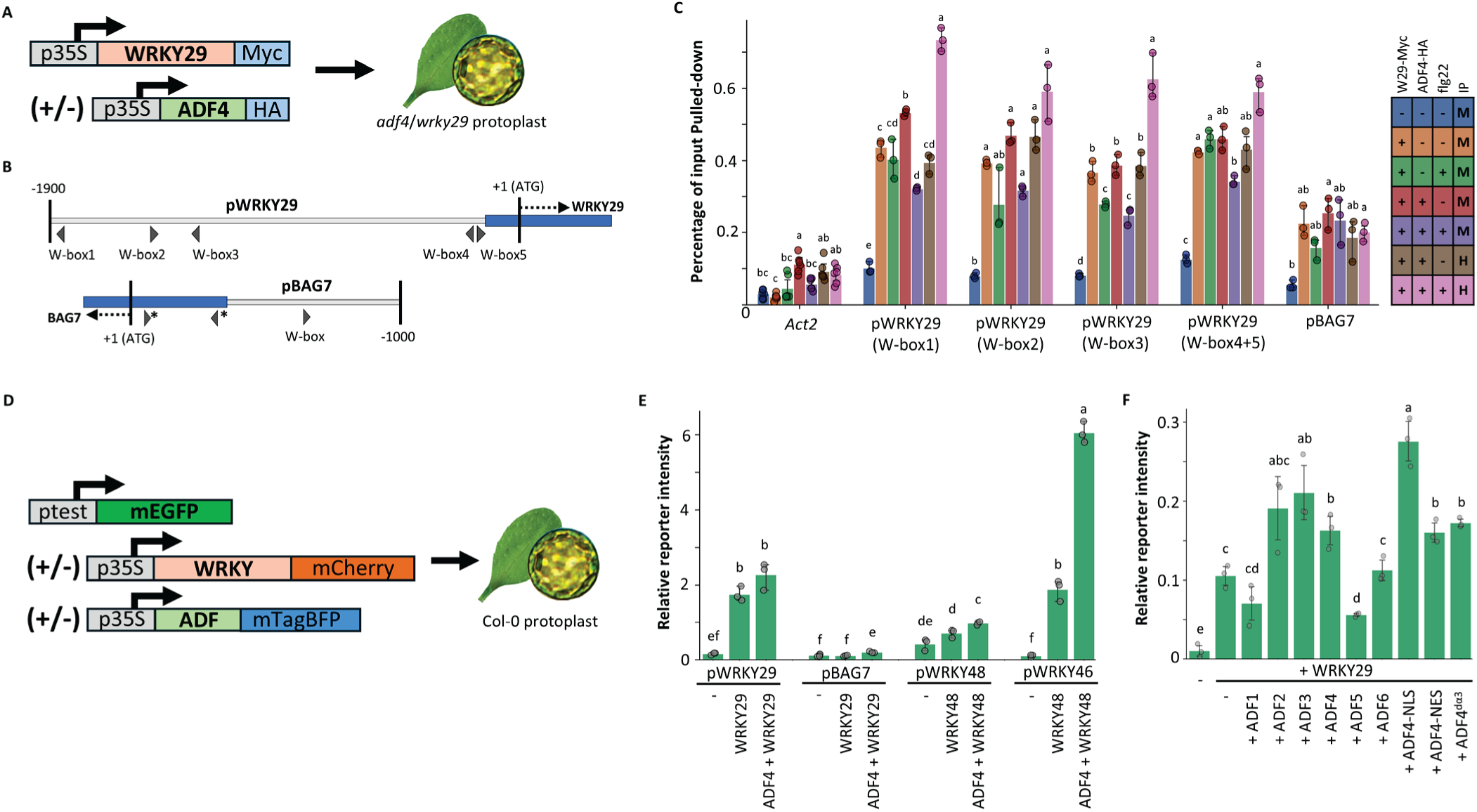
ADFs associate with WRKYs at targeted promoters and regulate gene expression level. **A**, experiment scheme of ADF4-WRKY29 ChIP assay. **B**, distribution of W-boxes on pWRKY29 and pBAG7. **C**, ChIP-qPCR quantification of W-box fragments enriched by WRKY29 and ADF4. WRKY29 and ADF4 (when co-expressed with WRKY29) can enrich pWRKY29 and pBAG7 fragments. *ACT2* gene serves as the negative control for enrichment quantification. flg22 treatment for 2h does not significantly impact the binding affinity of WRKY29 and ADF4. **D**, experiment scheme of the promoter reporter assay measuring ADF activities. **E**, promoter reporter assay of WRKY29/48 on different promoters. **F**, promoter reporter assay testing the impact of different ADFs on WRKY29-targeted pWRKY29. All data pass ANOVA with P<0.05; non-overlapping alphabets suggest P<0.05 upon post-hoc T-tests adjusted by Benjamini–Hochberg procedure for family-wise error.

As shown in Fig. 4C, WRKY29-MYC was enriched in all W-box-containing fractions of pW29, suggesting WRKY29 indeed binds to the W-boxes of its own promoter. Interestingly, when WRKY29-Myc and ADF4-HA were co-expressed, both α-HA and α-Myc ChIP were able to significantly enrich the pW29 W-boxes, indicating that WRKY29 and ADF4 form a complex together with their targeted DNA. Next, we introduced the treatment of flg22 in the ChIP experiment. However, the impact of flg22 was not obvious, indicating PTI activation is not a prerequisite to trigger the ADF4-WRKY29-DNA interaction under our transient expression condition. Although pBAG7 was also enriched by WRKY29 and ADF4 ChIP, the enrichment was not as strong as that for pW29.

### ADF2/3/4 enhance the WRKY29-mediated transcriptional activation

Next, we investigated if the ADF-WRKY interaction leads to regulation of the target genes of WRKYs. To test this, we constructed a promoter reporter system to quantify the activity of WRKY in the presence/absence of ADFs. The reporter system is comprised of 3 vectors, which contains p35S::WRKY-BFP (mTagBFP), p35S::ADF-RFP (mCherry), and GFP (mEGFP) driven by WRKY-targeted promoter (see Fig. 4D). When the GFP reporter vector is co-transformed with a combination of WRKY-BFP and/or ADF-RFP into protoplasts, all three fluorescence signals can be quantitatively measured by confocal microscopy, and the levels of GFP fluorescence indicate transcriptional activation amplitudes by WRKYs and/or ADFs. pW29 and pBAG7 were tested for WRKY29, whereas pW48 (assumed as auto-regulated by WRKY48) and pW46 (Gao et al., 2013) were used to test WRKY48. The *wrky29* and *wrky48* mutants were used as the sources of protoplast to eliminate the endogenous background WRKYs. As shown in Fig. 4E, both WRKY29 and WRKY48 were able to boost the expression of their respective targeted promoters. More importantly, ADF4 further enhanced the transcriptional activation amplitude when co-transformed with WRKY29.

We continued to test other leaf-expressed ADFs (ADF1-6) for their potential effects on WRKY activity. Plus, we also included ADF4-NLS and ADF4-NES (<u>n</u>uclear <u>e</u>xport <u>s</u>ignal) to evaluate the contributions of ADF4 as a function of its nuclear and cytoplasmic localization. As shown in Fig. 4F, ADF2/3/4 boosted the activity of WRKY29, while ADF1 and ADF5 did not. ADF4-NLS led to the highest transcriptional activity of pW29, which supports the hypothesis that the nuclear localization of ADF4 is important to enhance WRKY29 activity at the targeted promoters. Of note, ADF4-NES also showed a relatively high reporter signal, similar to that observed for ADF4. Further computational analysis showed that this was potentially caused by fluctuated ADF/WRKY expression ratios inherent to combinations of different test groups. In tune with this hypothesis, the normalized effect of per unit of ADF4-NES was much lower compared to ADF4 (supplemental Fig. 6). Additionally, ADF4^dα3^ was enhanced the activity of WRKY29 to a level similar to ADF4 (Fig. 4F), confirming that the ability of ADF4 to boost WRKY transcriptional activity does not rigorously require its interaction with actin. The sum of these data mirrored the BiFC results (Fig. 3E), showing that strong WRKY interactors – ADF2/3/4 – enhance the transcriptional activity of WRKY29 and that, at least in the case of ADF4, this enhancement primarily occurs inside the nucleus.

### ADF4, binding WRKY29, targets immune genes genome-widely and regulates WRKY29 targeting affinity and spectrum

Since our study revealed ADFs as a WRKY regulator, our next major question is how ADF’s WRKY binding and regulatory activity can contribute to plant immunity. Therefore, we conducted a set of ChIP-seq experiments to learn what the function of WRKY29 is as a TF and how ADF4 is involved when forming the complex. In brief, we transformed different combinations ADF4-HA and WRKY29-Myc to *wrky29/adf4* protoplast to pull-down ADF- and WRKY-bound genome fractions (Fig. 5A). A list of promoter fraction enrichment levels of all samples is provided as Supplemental Material 2.

**Figure 5:**
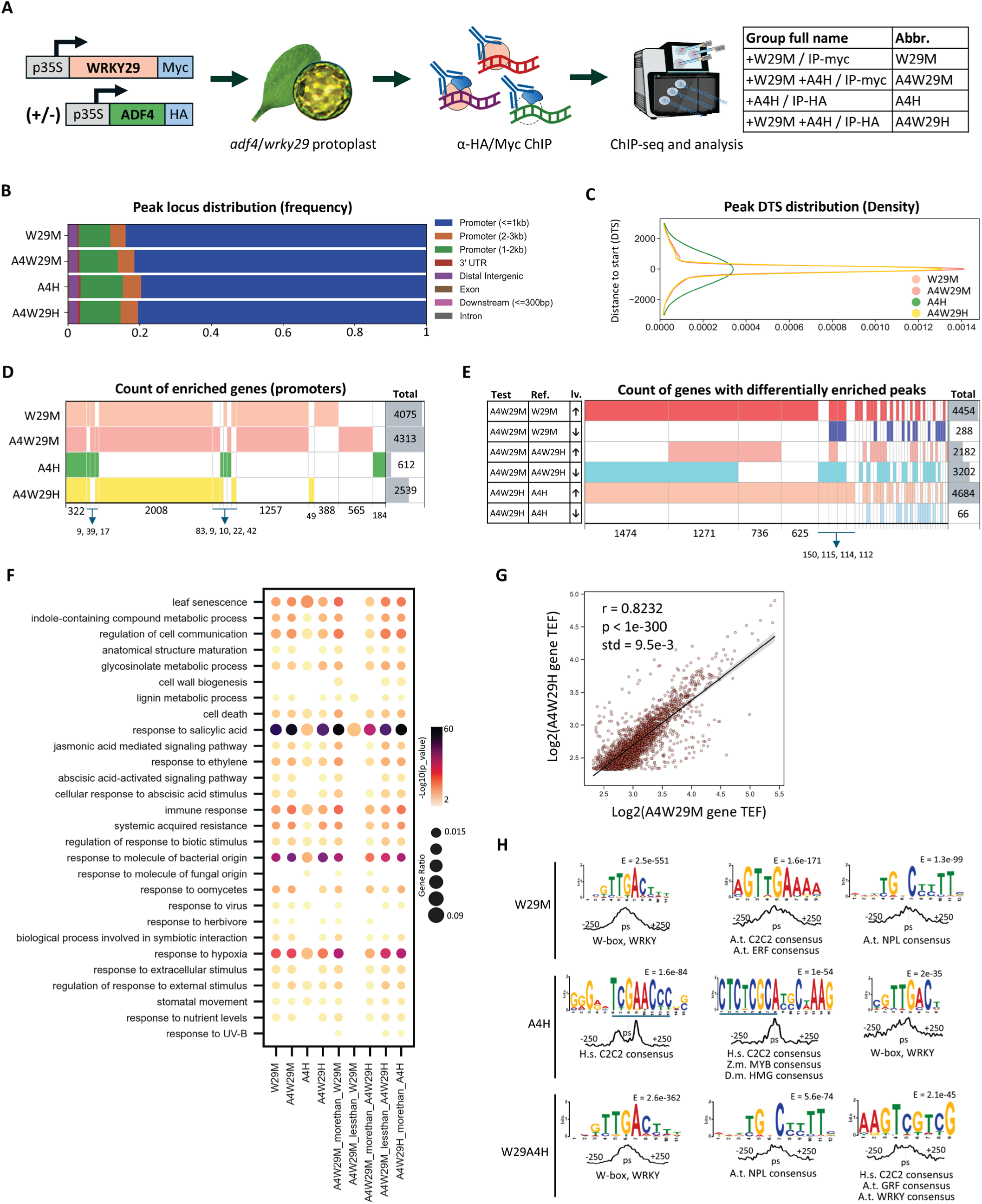
ChIP-seq reveals genome-wide immune gene promoters co-targeted by WRKY29 and ADF4. **A**, the ChIP-seq scheme. Experiment group abbreviations are described. **B**, Peak distribution of pulled-down genome fragments, by genomic function. Regardless of experiment groups, most peaks are located on promoters. **C**, The distance-to-start (DTS) distribution of peaks. **D**, SuperVenn diagram describing the overlapping or specificity levels of enriched gene promoters among different groups. **E**, SuperVenn diagram describing the overlapping or specificity levels of differentially enriched single peaks among different comparisons. **F**, GO pathway enrichment analysis on genes promoters enriched by ChIP and those with differentially enriched single peaks. Selected pathways are presented. **G**, Correlation analysis of gene enrichment levels by WRKY and ADF. Scatter plot represents the logarithmic enrichment level (defined by TEF, see Methods) of gene promoters identified by both A4W29M and A4W29H. A strong linear correlation with r = 0.8232 is detected, suggesting WRKY29 and ADF4 colocalize on their genome targeting sites. **H**, cis-element enrichment analysis identifies motif consensuses by binding sites of WRKY and TF in other families. MEME-ChIP were used analyze the ±250bp region surrounding the summit of each peak in different groups. TF binding consensus of WRKY, C2C2, ERF, NPL, MYB, and HMG family are identified. Blue bar highlights nucleotides when only partial of the motif is used to search for TF binding consensus.

The ChIP-seq confirmed that WRKY29 mostly binds to the promoter regions like a typical TF (Fig. 5B, C). A total of 4075 genes covering various immune pathways are targeted by WRKY29 (group name: W29M, Fig. 5D, F), providing the first comprehensive inventory of the targeting spectrum of WRKY29 *in planta*. The high number of target promoters indicates that WRKY29 is likely a broad-spectrum immune-regulatory TF. Surprisingly, when ADF4 was transformed alone (group name: A4H), it still targeted 612 genes that partially overlap with those of WRKY29 (Fig. 5D) and was also localized mainly on promoters (Fig. 5B, C). With respect to the biological function of the target promoters, A4H resembles the pattern of W29M, which is enriched with SA-immunity related genes, although the enrichment level and significance were not as high as W29M (Fig. 5F). Given that ADF4 does not have any DNA-binding domain, it is likely that ADF4 binds to other TFs or other components of the transcriptional machinery as a broad-spectrum transcriptional regulator.

The most interesting results were observed when ADF4 and WRKY29 were co-transformed. For WRKY29 (group name: A4W29M), ADF4 changed its targeting spectrum, as additional 687 new promoter loci were found and 449 WRKY29 targets got lost (W29M vs A4W29M, Fig. 5D). In addition, ADF4 significantly increased the enrichment level of single peaks covering 4454 promoters while decreasing those of 288 promoters (Fig. 5E), suggesting that ADF4 substantially enhances or changes promoter binding of WRKY29.

For ADF4 (group name: A4W29H), on the other hand, the addition of WRKY29 greatly changed its targeting spectrum. First, WRKY29 enables ADF4 to interact with additional 2191 gene promoters (Fig. 5D), with peaks of ∼4700 genes showing significantly changed (mostly upregulated) enrichment level (Fig. 5E). Most of the enriched genes by A4W29H overlap with those by A4W29M/W29M (Fig. 5D) and share the same function (Fig. 5F). Since WRKY29, rather than ADF4, has the direct DNA binding structure, the acquired DNA binding spectrum of ADF4 most likely involves the formation of the ADF4-WRKY29 complex. To further inspect this hypothesis, we performed a co-localization assay using the enrichment fold of overlapped genes from A4W29M and A4W29H (Fig. 5G). As A4W29W and A4W29H showed strong and significant linear correlation, showing that the ADF4-WRKY29 complex spread over the promoter loci across the genome. Combining previous results, we concluded that ADF4, through its physical interaction with WRKY29, can actively exert a substantial genome-wide regulatory effect on the immune transcriptome.

Next, we conducted a *de novo* motif enrichment analysis using the peak-summit neighboring sequences to interrogate the biochemical features of the WRKY and ADF-WRKY complex from a more comprehensive perspective (Fig. 5H). As expected, we found that WRKY29 utilizes W-boxes as its theoretically targeted motif. Besides, WRKY29 may also integrate into larger complexes with other TFs, since the binding motifs of other DNA binding protein families were enriched as well. For ADF4 without WRKY29, the enriched motifs reflected various families including C2C2 (family of *Hs*ZMYND8, a target of *Hs*Drebrin), MYB, HMG, and WRKY, suggesting that ADF may extensively regulate other DNA-binding proteins beyond WRKYs. By introducing WRKY29 to ADF4, the motif pattern enriched by ADF4 revert back to a pattern resembling that of WRKY29, echoing our previous conclusion that WRKY29 guilds ADF4 to its targeting spectrum to co-regulate the immune transcriptome.

### Nuclear ADFs have specific, pathogen-related roles in plant immunity

Considering the observed function of nuclear ADF, we are curious about whether and to what extent nuclear ADFs contribute to plant immunity. To address this question, we constructed complementation lines expressing mTagBFP fusions of ADF4 or ADF4-NES/NLS in the *a234* mutant background to restrict ADF4 within the cytosol or the nucleus (Fig. 6A). As expected, ADF4-NES displayed typical actin bundle localization in the cytosol without nuclear signal, whereas ADF4-NLS was restricted within the nucleus. The native ADF4 protein was observed in both the cytosol and the nucleus (Fig. 6B, C).

**Figure 6:**
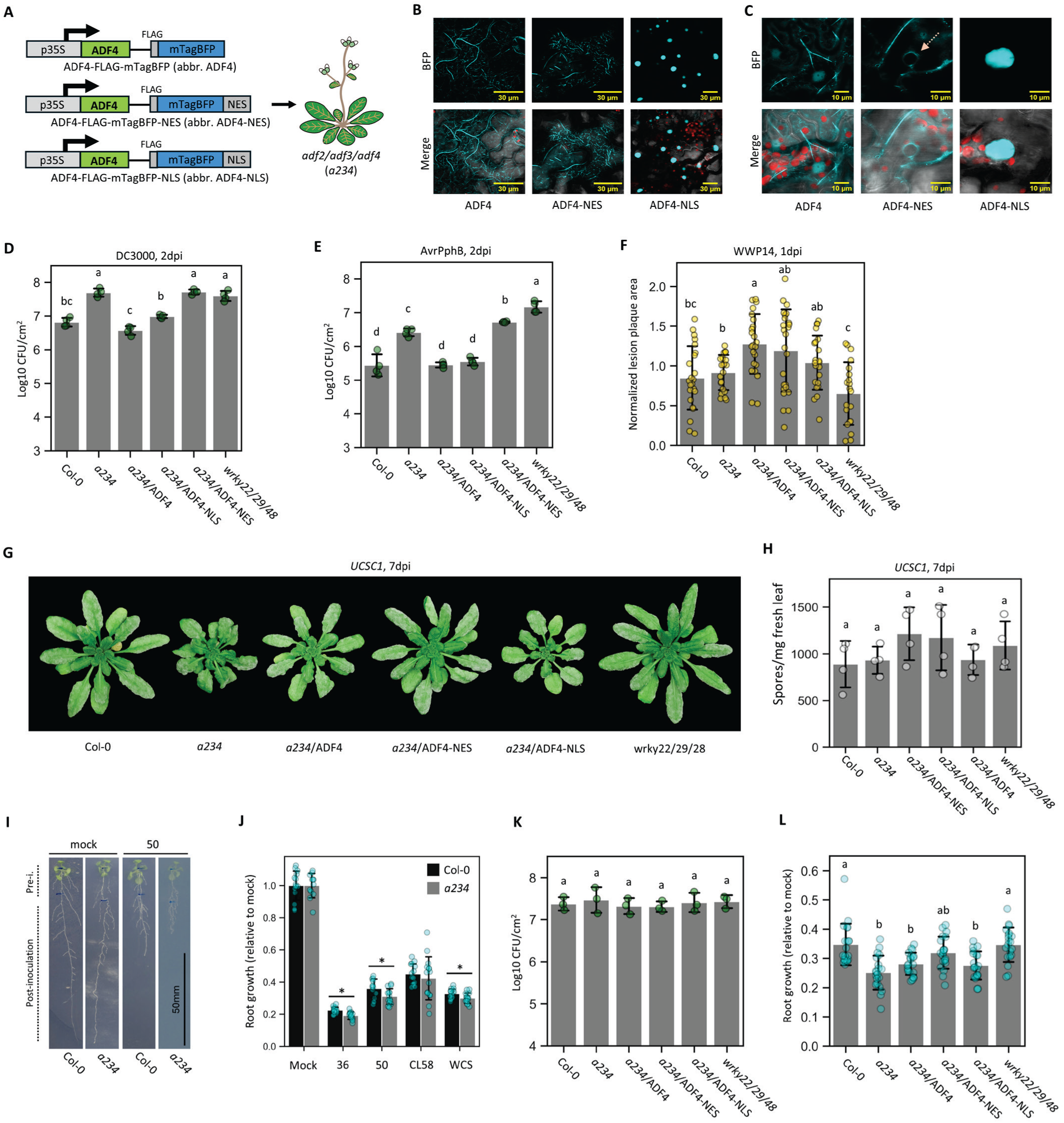
cytosolic and nuclear ADF4 have distinguished roles in different scenarios of plant immunity. **A**, Scheme of the construction of general and cytoplast/nucleus-exclusive ADF complementation lines. **B**-**C**, subcellular localization of ADF4, ADF4-NES (nuclear exiting signal), and ADF-NLS (nuclear localization signal) in Arabidopsis, displayed by low (**B**) and hgh (**C**) magnification. ADF4 normally localizes in both cytosols, with aggregation on actin bundle, and nucleus. NES and NLS completely forced cytosolic and nuclear localization. Cyan = ADF4 (mTagBFP); red = chloroplast; gray = bright field. Pink dashed arrow points to an empty nucleus. **D**-**E**, *P. syringae* growth essay on different *a234* and ADF complementation lines, using DC3000 (**D**), and DC3000/AvrPphB (**E**). Bacteria was inoculated to 5-week-old plants at OD600 = 0.002. Samples were harvested at 2dpi. See supplemental Fig. 8 for 0dpi data. Dampened resistance in *a234* can be totally rescued by ADF4 or majorly by ADF4-NLS, but not ADF4-NES. wr*ky22/29/48* phenocopies *a234*. **F**, Pectinobacteria growth essay on different ADF complementation lines. Col-0, *a234*, and *w22/29/48* show no difference among each other, but overexpressing ADF4 increases susceptibility. **G**-**H**, Powdery mildew *G.c.* UCSC1 infection assay. No significant difference among genotypes was detected by visual observation (**G**) or conidia spore quantification (**H**). **I**, demonstration picture of root commensal bacterial infection vertical plate assay (VPA). 1-week-old axenic seedlings are carefully transferred to a vertical plate previously spread with root commensal bacteria, root architectures are imaged 10 dpi to determine the developmental impact by bacteria. **J**, root growth VPA upon commensal bacterial using *a234*. Bacteria 36, 50, and WCS (see strain details in Methods) have enhanced growth inhibition on *a234*. **K**, bacteria growth in VPA on *a234* and complementation lines inoculated with 50. 50 do not show different population among Arabidopsis genotypes. **L**, Root growth in VPA on *a234* and the complementation lines inoculated with 50. *a234* has enhanced root growth inhibition but cannot be complemented with ADF4 or ADF4-NES/NLS. *wrky22/29/48* does not phenocopies *a234*. Data of **D**, **E**, **F**, **J**, and **L** pass ANOVA with P<0.05; non-overlapping alphabets suggest P<0.05 upon post-hoc T-tests adjusted by Benjamini– Hochberg procedure for family-wise error.

To investigate to which extent different localizations of ADF4 contribute to plant immunity, we first tested the respective transgenic lines for their resistance against *P. syringae*. As shown in Fig. 6D (0dpi: Supplemental Fig. 7), *a234*, which phenocopies *wrky22/29/48*, has dampened basal defense against DC3000. The *a234* phenotype was fully complemented by expressing native *ADF4*, partially complemented by ADF4-NLS, but not by ADF4-NES. Further, the resistance levels seemed to be positively corelated to the expression levels of ADF4/ADF4-NLS among the transgenic lines (Supplemental Fig. 8). When DC3000/AvrPphB was used to trigger ETI, the major profile of resistance was similar to DC3000, and ADF4-NLS fully restored resistance (Fig. 6E). These results suggested that nuclear – rather than cytosolic – ADF drives the majority of the resistance against DC3000, which is even more critical upon AvrPphB-induced ETI.

Next, we interrogate whether the immune function of ADF4 – especially its nuclear fractions – is general or pathogen-specific. Particularly, we chose *Pectobacterium carotovorum*, the pathogen of bacterial soft rot, as a representative of absolute necrotrophs and *Golovinomyces cichoracearum, the* fungal pathogen of powdery mildew, as a representative of absolute biotrophs, for pathogen growth assay. When *P. carotovorum* WWP14 was inoculated, we did not observe significant difference between Col-0, *a234*, and *wrky22/29/48*. (Fig. 6F). However, lines overexpressing ADF4 are more susceptible to WWP14, indicating a potential dosage-dependent role of ADFs in this defense process. For powdery mildew, there is no significant difference in the fungal growth among genotypes inoculated with *G. cichoracearum* UCSC1 (Fig. 6G, H). This observation shows an interesting discrepancy from the previous study that suggested an enhanced resistance against powdery mildew in *ADF1/2/3/4*-silenced line (Inada et al. 2016). However, it is noteworthy that their study uses a different powdery mildew species, *G. orontii*, which may explain the different phenotypes. Inspired by this hypothesis, we tested the *G. cichoracearum* UCSC1 growth on the same *ADF1/2/3/4*-silenced line, and rather than resistance, we observed contrast greater susceptibility compared to Col-0 (Supplemental Fig. 9), which suggests that ADF’s roles in pathogenesis of different powdery mildew species can be substantially different.

To understand the potential reasons why the immune function of (nuclear) ADFs is pathogen specific, we conducted a comprehensive analysis on 51 publicly released transcriptome datasets (Yu et al. 2022; Li and Xiao. Accepted.) describing Arabidopsis challenged by *P. syringae*, as well as other biotrophs and necrotrophs (Supplemental Fig. 10). We found that Arabidopsis has varied gene regulatory profiles when infected by different pathogen species, in a degree of contrast greater than those comparing basal defense vs ETI (Supplemental Fig. 10A, B). Specifically, the transcriptome profile of DC3000 infection is dramatic different from *G. cichoracearum* infection (Supplementa Fig. 10C) and moderately different from *Botrytis cinerea* (a necrotroph generating similar symptoms of *P. carotovorum*) infection (Supplemental Fig. 10D). The genes attributable to this discrepancy are generally involved in metabolism of growth/defense molecules, hormone signaling, and defense and other stress response (Supplemental Fig. 10E). However, there are only ∼32% and ∼21% of those genes also regulated by ADF2/3/4 in the individual cases of *G. cichoracearum* and *B. cinerea*, respectively (Supplemental Fig. 10F, G; ADF-regulated genes are described later). Therefore, we infer that the differences in pathogenesis of these species, causing different transcriptome regulatory profiles, provide a potential explanation for the different disease phenotypes related to nuclear ADFs.

In addition to typical pathogens, we also investigated whether nuclear ADF influences the interaction with root commensal bacteria as a perspective from wide concept of plant-microbe interaction. Such bacteria may proliferate in root apoplast and regulate host morphogenesis and immunity without causing disease. We started by measuring root growth upon different commensal bacteria treatment, as an index reflecting host response commensal microbes (Fig. 6I). As a result, we identified that *a234* showed a minor root zig-zag morphology and a more severe root growth inhibition upon mono-inoculation with several species of commensals (Fig. 6J). Regarding this, we chose Arabidopsis rhizosphere *Pseudomonas sp.* KD5 (IMG ID: 2228664007, lab internal ID: “50”) as a representative strain for further analysis. When “50” was inoculated onto different genotypes, there was no significant difference over bacterial growth rate (Fig. 6K). However, *a234* displayed stronger root growth inhibition following “50” treatment, which was not fully complemented by ADF4 and its localization variants. Meanwhile, *wrky22/29/48* did not phenocopy *a234*, suggesting these WRKYs are not critically involved in root growth regulation by rhizosphere commensal bacteria. Combining current knowledge (García-González and van Gelderen 2021), we assume that ADF2/3, rather than ADF4, majorly contribute to root development in an actin-related manner that resembles the roles of ADF4 in hypocotyl development (Yu et al. 2025), which is dominated by its cytosolic function.

In summary, the immune function of ADFs varies among different types of pathogens. Nuclear – but not cytosolic – ADFs contribute to basal defense and at least certain types of ETI against *P. syringae*. However, they do not impact defense against *P. carotovorum* WWP14 or *G. cichoracearum* UCSC1 at natural expression level, but overexpressed ADF4 can enhance WWP14 susceptibility. Besides, ADFs are also involved in microbe-regulated root morphogenesis. These results indicate that the cytoplasmic-nuclear function of ADF is relatively complex across different types of plant-microbe interaction.

### Nuclear ADFs regulate transcriptome at the resting state and immune events

Since nuclear ADFs contribute to plant immunity and interact with transcription machinery, we conducted a series of mRNA-seq experiments to further dissect the role of nuclear ADFs on plant immunity. RNA samples from Col-0, *a234*, and ADF4/NES/NLS complementation lines, with mock or PAMP (flg22) treatment, were collected and sequenced to quantify the transcriptome at the resting and PTI-activated states, respectively (Fig. 7). Overall, both PAMP elicitation status and genotype contribute to the variation of transcriptional profile (Fig. 7A-C, Supplemental Fig. 11, 12). We totally identified 4389 deferentially expressed genes (DEGs) related to genotypes (Supplemental material 3); principle component analysis (PCA) suggested that Col-0 vs *a234* showed the largest difference, which can be complemented by ADF4 or ADF4-NLS, but not ADF4-NES (Fig. 7C), revealing the general impact of nuclear ADFs on the immune transcriptome.

**Figure 7:**
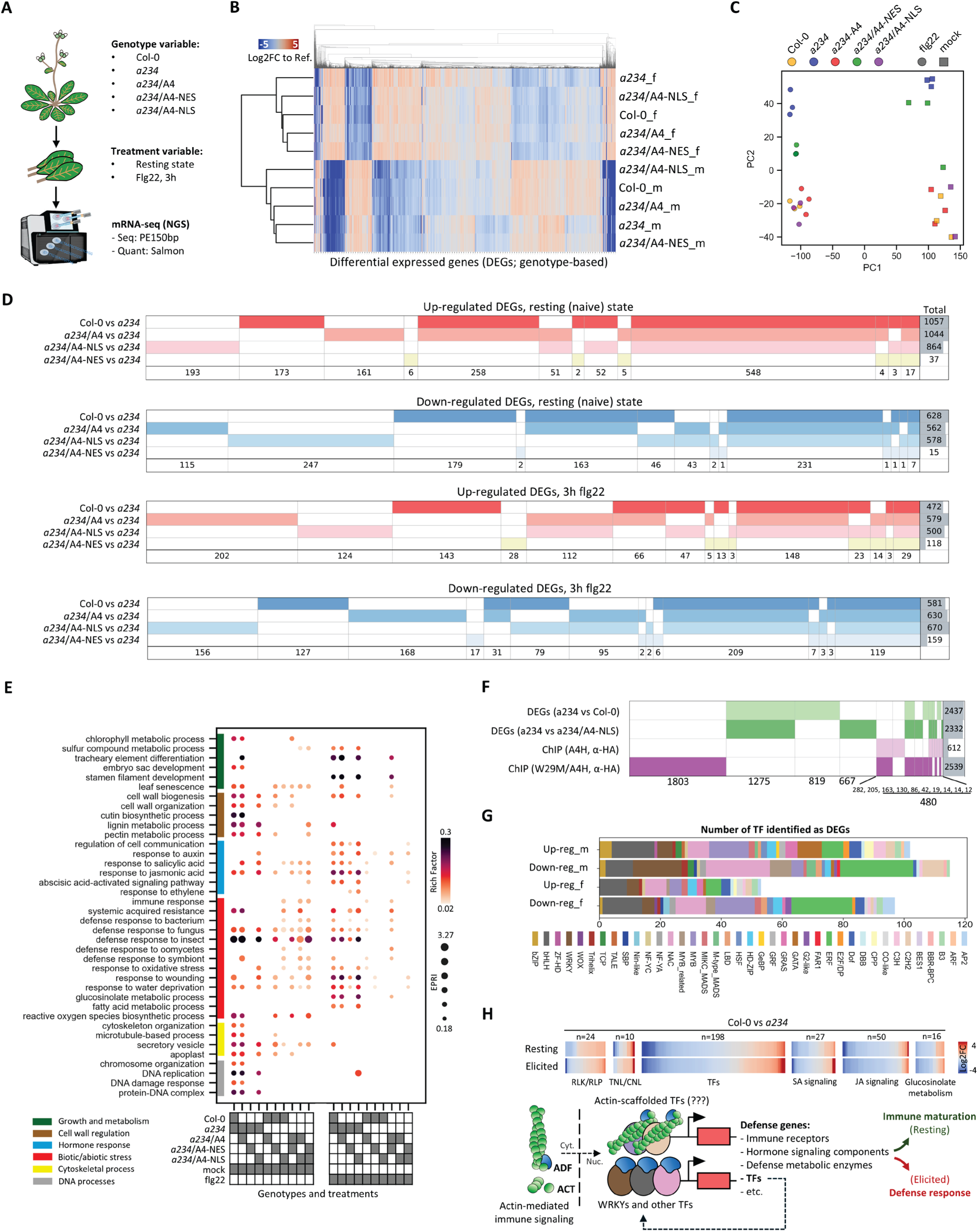
Nuclear ADF4 mediates transcriptomic regulation at resting state and PTI event. **A**, mRNA-seq experiment scheme. **B**, DEG clustering analysis. Heatmap represents the logarithmic fold-of-change of TPM compared to the respective mean expression level of all groups, which is clustered by their cross-group patterns. DEG is identified by DEseq2, defined by significant deference of FPKM across any two genotypes at the same treatment conditions (see Methods for detail). **C**. 2D-PCA of all samples. ADF4 and ADF4-NLS successfully complement *a234* at both resting and elicited status. **D**, SuperVenn diagram of overlapped and specific DEGs regulated by ADF2/3/4. For easy visualization, data are split by two variables: resting state (top 2) and PTI event (bottom 2), as well as up-regulation (red) and down-regulation (blue). For all conditions, normal ADF4 complements *a234*, ADF-NLS majorly complements *a234* with a few extra DEGs, but ADF4-NES cannot complement *a234*. For the complete and all-in-one SuperVenn, please refer to Supplemental Fig. 12. **E**, GO pathway enrichment analysis on genes DEGs among different groups. Selected pathways related to growth and metabolism, cell wall regulation, hormone, stresses (including immunity), cytoskeleton, and DNA processes are presented. We use Euclidean Pathway Regulation Index (EPRI), a normalized logarithmic indicator to measure the level pathway regulation by DEGs (see *Methods* for reference). ADF4 and ADF4-NLS complementation mimic the transcription regulatory function of ADF2/3/4 in Col-0, while ADF4-NES does not. DEGs in mock and flg22-treated samples exhibit different patterns, suggesting partially distinguished role of nuclear ADF4 in the different phases. **F**, SuperVenn diagram comparing ADF4-regulated genes and ChIP-identified ADF4-targeted gene promoters. A sum of 480 genes is both regulated and physically targeted by ADF4, as underlined. **G**, Distribution of ADF4-regulated TFs by protein family. ADF4-regulated genes are defined by the union set of DEGs of Col-0, *a234*/A4, or *a234*/A4-NLS vs *a234* in any condition. **G**, Summary of ADF-mediated transcriptional regulation. Heatmaps describe the ADF regulatory function on selected groups of defense-related DEGs, as a demonstration of model concept.

Our investigation into the distribution and functionality of the DEGs led to three major discoveries. First, compared to *a234*, Col-0 has ∼1700 DEGs at the resting state (i.e., mock treatment) and ∼1000 DEGs upon PTI activation (Fig. 7D, supp. Fig. 13). These genes reflect various physiological processes including immunity, metabolism, hormone signaling, cell wall regulation, cytoskeleton organization, and DNA processes (Fig. 7E). Notably, *a234*/A4 largely phenocopies Col-0 when they are compared with *a234*. Therefore, ADF2/3/4 indeed significantly contributes to the expression of defense and other genes and function redundantly in gene regulation. Second, ADF4-NLS complementation dramatically changed the transcriptome pattern of *a234* by ∼1500 DEGs over resting state and ∼1200 DEGs upon PTI, whereas *a234*/A4-NES was almost identical to *a234* (Fig. 7D, Supplemental Fig. 13), which, echoing results above, validated that nuclear – not cytosolic – ADFs possess the major transcriptional regulatory activity *in planta*. Third, nuclear ADFs regulate different genes and biological pathways at the resting and elicited status. While (mostly nuclear) ADFs are involved in immune regulation in both naive and PTI-triggered plants (Fig. 7C, D), genes related to cell wall regulation, cytoskeleton regulation, and DNA processes are exclusively and heavily regulated by ADFs in the naive plant (Fig. 7E). On the other hand, when PTI is triggered, DEGs shift to immune-related glucosinolate and fatty acid metabolic processes, with extended spectrum of hormone regulation (auxin, ABA, and ethylene, beyond SA and JA; Fig. 7E). Therefore, nuclear ADFs regulate different groups of genes at the nexus of establishing the growth-defense equilibrium.

Regarding the ADF-WRKY coregulatory mechanism revealed by this study, we proceeded to inquire whether – or to what extent – the promoter binding activity of the ADF4-WRKY29 complex could explain the dampened immune transcriptome due to the lack of ADFs. We compared all DEGs revealed from between col-0, *a234*, and *a234*/A4-NLS, with ADF4 targeted genes identified through ChIP (Fig. 7F). We found 480 genes that are both regulated by ADFs at expression level and physically targeted by ADF4 on their promoters. However, the majority of the DEGs or ADF4 targeted genes do not overlap, indicating that the co-regulation on promoters by the ADF-WRKY complex – at least by merely ADF4 and WRKY29 – may not fully explain the transcriptional functionality of ADFs. As a further annotation to the nuclear activity of ADFs, our categorical analysis suggested that ADFs regulates the expression level of 309 TFs, covering 39 TF families beyond WRKYs (Fig. 7G). Considering that nuclear ADF4 has specific affinity to 67 TFs through TurboID (Fig. 2H), we propose that ADFs mediate a broad spectrum of transcriptional regulation potentially by directly interacting with certain TFs to regulate their activity, which influences the expression level of additional TFs through their inter-regulatory network, thereby facilitating broad immune impact. While this activity may not require the actin-ADF interaction, we cannot totally exclude the possibility that ADF’s nuclear function may be indirectly affected by its actin interaction under other conditions. Therefore, nuclear ADFs potentially may have multiple layers of function regarding transcriptional regulation, where those related to the ADF-WRKY complex represent a prominent example, which has been revealed in this study.

## Discussion

Plant immune activation mobilizes the cytoskeleton, including its associated proteins, to transport and stabilize defense molecules and organelle, as well as numerous additional physiological responses (e.g., pathogen triggered stomatal closure). Due to the significant roles of cytoskeleton remodeling in plant immunity, it is reasonable to deduce that critical proteins directly regulating cytoskeletal dynamics are indispensable for plants to render full strength of immune response. Massive experimental evidence matches this theory (Li and Day 2019), making it a popular mechanism model to explain why actin remodelers, such as ADFs, influence pathogen susceptibility and resistance. However, since this process is theoretically happening in the cytosol, the canonic function of ADF cannot explain why ADF evolved to bring actin into nucleus (Dopie et al. 2012). Moreover, it is unclear how nuclear-localized ADFs contributes to certain types of resistance (Inada et al. 2016), which implies a different type of moonlighting function currently undescribed.

In this study, we used the CRISPR-Cas system to eliminate the redundancy of Arabidopsis leaf-expressed ADFs, successfully defining the nuclear-exclusive function of ADF4, as a representative of Class I ADF, in plant immunity. As an explanatory mechanism, we identified WRKY22/29/48 as strong interactors of ADF2/3/4, providing evidence that these interactions regulate the promoter activity and targeting spectrum of WRKYs. Through this study, we propose a novel perspective to review the role of ADF proteins in plant immunity and beyond – a function that is relatively independent from its traditional identity as a regulator of actin remodeling. Here, we would like to discuss the significance, potential, uncertainty, and limitations of the newly identified transcriptional regulatory function of nuclear ADFs.

As illustrated in Fig. 8, we summarized the discoveries and supported hypotheses for an integrated cytoplasmic-nuclear functional model of actin and ADF. Our study, combined with previous work in this field, notes that a full and robust immune system requires ADF to perform both actin severing/depolymerization activity in cytosol and transcriptional regulation activity in the nucleus, integratively. As seemingly straightforward as this activity may appear, this integrated dual-phase function is more complex. As previously noted, immune signaling can activate upstream kinases capable of phosphorylating corresponding ADFs (Zhao et al. 2016; Lu et al. 2020). When ADF is phosphorylated (e.g., S6 and S105 of ADF4, or S3 of *Hs*CFL), it will dissociate from actin, thus eliminating the traditional role of ADF as a actin depolymerization factor, which is regarded as a central mechanism to regulate actin architecture, and including immune-triggered actin remodeling. As presented herein, ADF-ACT and ADF-WRKY interactions do not share interfaces on ADF (Fig 3E, F; Supplemental Fig. 4), so we posit that ADF phosphorylation events decrease the ADF-ACT interaction intensity and hypothetically increase the ADF-WRKY interaction intensity. At present, we still do not know if phosphor-regulation either directly enhances the binding affinity between ADF and WRKYs, or if this is an indirect influence whereby phosphorylation promotes ADFs disassociation from ACTs, which in turn drives the ADF competition equilibrium to ADF-WRKY. If any of these mechanisms are valid, it indicates that ADF-ACT and ADF-WRKY interaction reflect two potential antagonistic aspects of ADF function. To fully test this hypothesis, a detailed dynamic phosphorylation profile of ADF during immune process *in vivo* is necessary, as well as a comprehensive analysis of ADF nuclear shuttling speed of different phosphorylation variants. We assert that these are critical questions to be addressed to fully define the ADF phospho-switch hypothesis.

**Figure 8:**
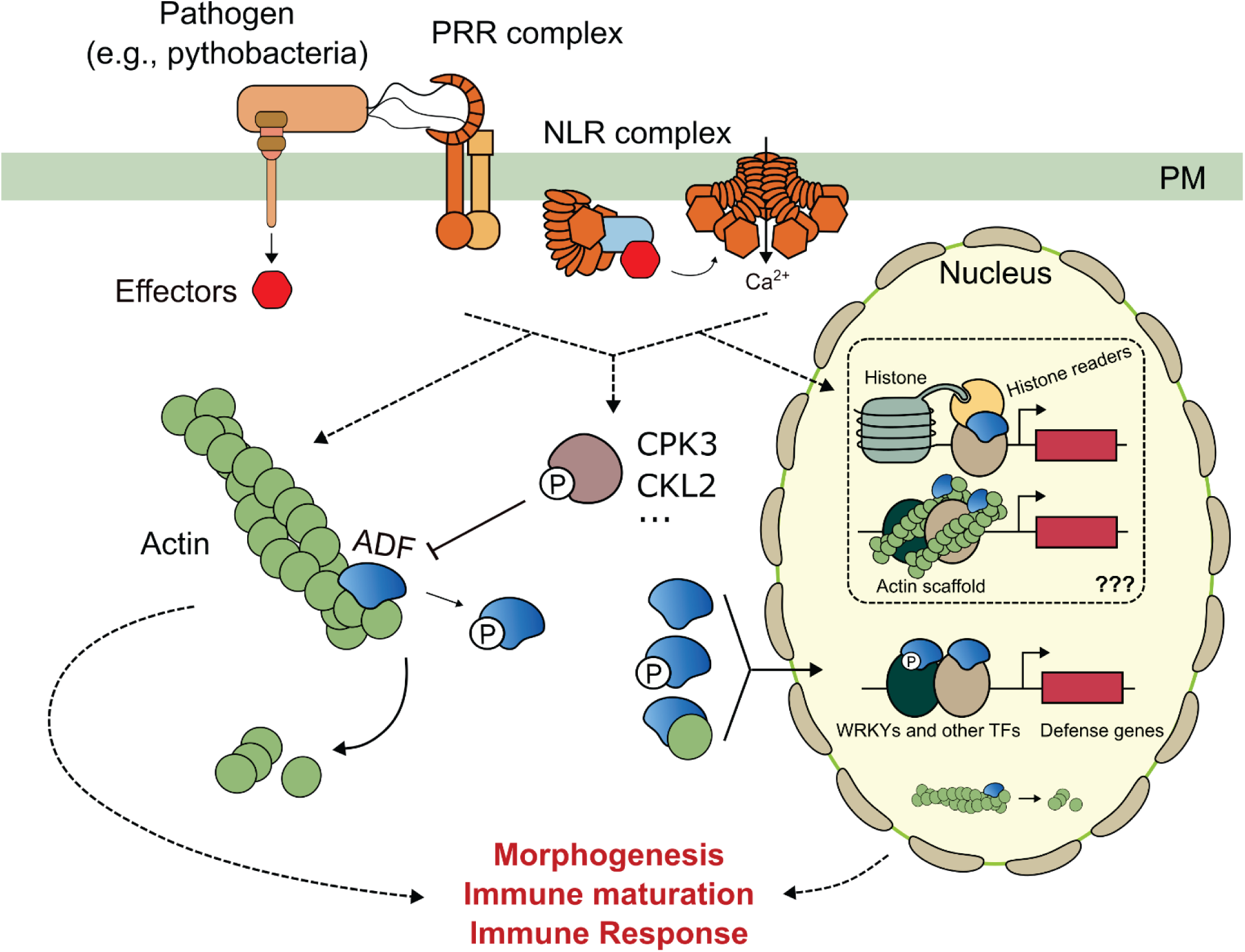
Proposed functional model describing the nuclear-cytosolic regulation of plant immunity by ADF. ADFs are duel-functional proteins that both (1) proceeding actin depolymerization, severing, or bundling in cytosol and (2) regulate transcriptional machinery in nucleus, as a moonlighting activity. These two activities are likely to utilize different protein-protein interaction interfaces on ADFs. Upon immune activation, ADF is phosphorylated by kinases such as CPK3 and CKL2, which blocks the ADF-actin interface and prohibits its actin depolymerization activity and changes the actin dynamics, thereby remodeling the actin cytoskeleton to a pro-immune phase. On the other hand, ADFs interact with the transcriptional machinery in the nucleus, including transcription factors (such as WRKYs) and presumably histone complexes as well as nuclear actin scaffolds, which regulates the defense genes and multiple aspects within and beyond immunity. Phosphorylation on ADF upon immune activation may enhance the ADF nuclear function by either directly enhancing its binding affinity toward WRKYs or indirectly providing more nuclear ADFs dissociated from cytosolic ACT to drive the equilibrium to ADF-WRKY side. The combination of cytosolic and nuclear function of ADF synergistically shapes plant immunity (including immune maturation and immune response) and other aspects of plant development.

As illustrated in Fig. 8, we summarized the discoveries and supported hypotheses for an integrated cytoplasmic-nuclear functional model of actin and ADF. Our study, combined with previous work in this field, notes that a full/robust immune requires ADFs to have both the actin severing/depolymerization activity within cytosol and the transcriptional regulation activity in the nucleus, integratively. As aforementioned, immune signaling can activate upstream kinases capable of phosphorylating corresponding ADFs (Zhao et al. 2016; Lu et al. 2020). When ADF is phosphorylated (e.g., S6 and S105 of ADF4, or S3 of *Hs*CFL), it will dissociate from actin, thus eliminating the traditional role of ADF as the actin depolymerization factor, which is regarded as a central mechanism to regulate actin architecture, including immune-triggered actin remodeling. As presented herein, ADF-ACT and ADF-WRKY interactions do not share interfaces on ADFs (Fig 3E, F; Supplemental Fig. 4), so we posit that ADF phosphorylation events decrease the ADF-ACT interaction intensity and would hypothetically increase the ADF-WRKY interaction intensity. At present, we still do not know if phosphor-regulation either directly enhances the binding affinity between ADF and WRKYs, or if this is an indirect influence whereby phosphorylation promotes ADFs disassociation from ACTs, which in turn drives the ADF competition equilibrium to forming the ADF-WRKY complex. If any of these mechanisms are valid, it indicates that ADF-ACT and ADF-WRKY interactions reflect two potential antagonistic aspects of ADF function. To fully test this hypothesis, a detailed dynamic phosphorylation profile of ADF during the immune process *in vivo* would be necessary, as well as a comprehensive analysis of ADF nuclear shuttling speeds of different phosphorylation variants. These represent critical questions to be addressed in the future to fully define the ADF phospho-switch hypothesis.

ADF2/3/4 are critical for full-strength pro-immune transcription (Fig. 7) as well as basal defense and specific sources of ETI against DC3000 (Fig. 1E-H, 6D-E), a process in which the nuclear – rather than cytosolic – action of ADF is required. Combined with ADF2/3/4’s ability to interact with WRKY22/29/48 and the results showing that *wrky22/29/48* phenocopies *a234*, we conclude that the nuclear ADF-WRKY interaction can largely explain the immune phenotypes. However, as noted in the TurboID data, there were additional ADF4-interacting TFs from other families beyond WRKY, as well as histones, histone readers, and modifiers (Fig. 2H, J). Correspondingly, the motif enrichment analysis using ChIP-seq reads enriched by ADF4 also discovered many cis-element patterns not directly related to the WRKY family. Hence, we infer that nuclear ADFs are potentially involved in multiple intertwined fractions of the transcriptional regulatory machinery, and that the ADF-WRKY interaction may be just the tip of the iceberg in terms of nuclear ADFs in transcriptional regulation under different conditions.

It is noteworthy that the DEG analysis identified that nuclear ADF4 also controls biological pathways related to development, metabolism, and DNA-related processes during the non-elicited (i.e., in the absence of pathogen infection) resting state. Combined with recent studies suggesting that ADFs regulate nuclear organization (Matsumoto et al. 2023) and senescence (Matsumoto et al. 2024), it is reasonable to deduce that nuclear ADFs have more general transcriptional regulatory functions beyond immunity, potentially by regulating other TFs or histone-based epigenetics. At the same time, such a mechanism potentially has a broader scope of application beyond Arabidopsis – or even beyond plants. As the ADF/cofilin family is widespread across almost all eukaryotic kingdoms that uses actin in any cellular process (Chen and Day 2024), we hypothesize that ADFs may play similar roles in transcriptional regulation in other plant and animal species, including human. Currently, there are many clinical diseases associated with dysfunctional ADFs/cofilins that exhibit abnormal transcriptome pattern (Shehjar et al. 2024; Xing et al. 2024), but this is stereotypically considered as an impact of abnormal or pathological actin dynamics. On the other hand, for plant diseases on crop species, most evidence remains at genetic level (Li and Day 2019), leaving greater intellectual challenges to rationalize how genotypes related to ADFs contribute to phenotypes in pathogenesis and immunity at cellular and molecular level. However, as we proved that ADFs can mediate immune and general transcriptional regulation through interaction with TFs, which is not necessarily related to ADF-ACT interaction, the cytosolic and nuclear ADF functionality is conceptually deconstructed into two relatively independent, but continuously inter-communicating, signaling nodules. Hence, we hope this study can provide the community with a novel perspective to explore the roles of ADFs in plant and human disease, as well as issues in general biology.

## Materials and Methods

### Plant and microbe materials

All plants were grown in a BioChambers model FLX-37 walk-in growth chamber (BioChambers, Manitoba, Canada) at 20 °C with 12h light/12 dark with 60% relative humidity and a light intensity of 120 μmol photons m^−2^ s^−1^. Arabidopsis mutant *adf1* (Salk_144459), *adf3* (SALK_139265), *wkry22* (CS68112), *wrky29* (CS339192), and *wrky48* (SALK_066438) were purchased from ABRC and screened for homozygosity. *adf4-2* (SALK_121647) were generously provided by Dr. Yan Guo. We identified that *adf4-2* contains another insertion in *AT1G77500*, so the original *adf4-2* was further screened to eliminate *at1g77500*. To further clean other potential insertions, *adf4-2/AT1G77500(WT)* was then backcrossed to Col-0 to obtain a purified *adf4-2* line, namely *adf4-2b*. The construction strategy and genotyping of all *ADF* and *WRKY* high order mutants are described in Fig. 1D and/or Supplemental Fig. 1.

*Pseudomonas. syringae* pv. tomato DC3000 and its AvrPphB-containing strains are preserved stocks of the Day Lab, Michigan State University (MSU), as previously described (Lu et al. 2020). *Pectobacterium carotovorum* WWP14 was generously gifted by Dr. Greg Howe (MSU). *Golovinomyces cichoracearum* UCSC1 is historically preserved in the Xiao Lab, University of Maryland, as previously described (Wu et al. 2024). Arabidopsis root commensal bacterial strains are preserved stocks of the Lebeis Lab, MSU. Strain identities as follows. WCS417: *Pseudomonas simiae* sp., IMG ID 2585427642; 50: *Pseudomonas sp.* KD5, IMG ID 2228664007; CL58, *Pseudomonas umsongensis* UNC430C58Col, IMG ID 2556921015; 36: *Pseudomonas mandelii* 36MFCvi1.1, IMG ID 2521172653.

### Construction of plasmid vectors

TurboID, BiFC, ChIP, promoter assay, and *ADF4* complementation in this study are enabled by newly constructed plasmid vectors, described as Supplemental Material 4. Related primers are listed in Supplemental Material 5.

### CRISPR gene editing

gDNAs targeting *ADF2* and *ADF4* were designed using CRISPR-P, as described in Supplemental Material 5. gDNAs were inserted into CRISPR-Cas9 vector psgR-Cas9-At (Mao et al. 2013) followed by insertion into pCAMBIA1300. Resultant constructs were then transformed into Col-0 and *adf3*, to knock out *ADF2* and *ADF4*, respectively, generating *adf2* and *adf3/4*. T1 antibiotic-resistance positive plants with chimeric polymorphism on targeted loci were selected; T2 plants with homozygous null mutation on targeted loci and without CRISPR vector insertion were selected for further experiments.

### Co-immunoprecipitation

Co-immunoprecipitation (co-IP) assays were performed based on a protocol adapted from (Lu et al. 2020), with slight modifications. In brief, ten *N. benthamiana* leaf discs (1 cm diameter) were harvested and frozen into liquid nitrogen. Leaf lysates were prepared by grinding leaf discs in liquid nitrogen and then grinded with 5mL homogenization buffer (50 mM Tris, pH 7.4, 150 mM NaCl, 10% glycerol, 0.2% triton X-100, 1mM DTT, 1 x cOmplete™ Protease Inhibitor Cocktail (Sigma #04693116001), and 1mM PMSF). Homogenized samples were sonicated at 12% amplitude for a total of 30s using a needle sonicator to release nuclear proteins and then centrifuged at 4 °C at 18600g for 5 min. 4 mL of the supernatant in a microcentrifuge tube was slowly rotated at 4°C for 4h with Pierce™ Anti-DYKDDDDK Magnetic Agarose (Thermo #A36797) following the manufacturer’s instructions. Samples – associated with magnetic agarose – were washed using homogenization buffer for at least 3 times, 5 min each. For western blot analysis, enriched fractions - beads with pulled-down fractions are mildly denatured by incubation with 1x LDS Sample Buffer (Thermo #B0008) and 50mM DTT at 70°C for 10min.

### TurboID and LC/MS/MS analysis

Arabidopsis TurboID protocol was adapted from a previous study (Mair et al. 2019). Briefly, 10 Arabidopsis leaves were first injected with a mixture of 1 μM flg22 and 1 μM elf18 or water as mock. Three hours later, the leaves were injected with 50 μM biotin and treated for 1 hour before frozen in liquid nitrogen. Frozen leaves were processed through a typical co-IP procedure as described above, except that an additional PD-10 desalting column (Cytiva #17085101) was used to eliminate the residual biotin in the centrifuged leaf lysate supernatant following the official protocol. Pierce™ Streptavidin Magnetic Beads (Thermo #88816) is used for pull-down biotin-labeled proteins.

For proteomics, the streptavidin beads binding TurboID targeted proteome were washed by 50 mM ammonium biocarbonate for 3 times. Trypsin, in the same buffer, was then added to the beads at 5 ng/μL so that the beads were just submerged in digestion buffer and allowed to incubate at 37°C for 6 hours. The solution was acidified to 1% with trifluoroacetic acid and centrifuged at 14000g. Peptide supernatant was removed and concentrated by solid phase extraction using StageTips1. Purified peptides eluates were dried by vacuum centrifugation and frozen at -20°C or re-suspended in 2% acetonitrile/0.1%TFA to 20 μL.

An injection of 5 uL was automatically made using a Thermo EASYnLC 1200 onto a Thermo Acclaim PepMap RSLC 0.1 mm x 20 mm C18 trapping column and washed for 5 min with buffer A (99.9% water/0.1% formic acid). Bound peptides were then eluted over 35 min onto a Thermo Acclaim PepMap RSLC 0.075mm x 500mm resolving column with a gradient of 8% buffer B (80% acetonitrile/0.1% formic acid/19.9% water) to 40% buffer B in 24 min. After the gradient the column was washed with 90% buffer B for 10 min at a constant flow rate of 300nl/min. Column temperature was maintained at a constant temperature of 50°C using an integrated column oven (PRSO-V2, Sonation GmbH, Biberach, Germany). Eluted peptides were sprayed into a ThermoScientific Q-Exactive HF-X mass spectrometer using a FlexSpray spray ion source. Survey scans were taken in the Orbi trap (60000 resolution, determined at m/z 200) and the top 15 ions in each survey scan are then subjected to automatic higher energy collision induced dissociation (HCD) with fragment spectra acquired at a resolution of 15,000.

### Bacterial and fungal pathogen growth assay

*P. syringae* inoculation and growth measurement was previously described (Lu et al. 2020). Briefly, DC3000 or DC3000/AvrPphB in 10 mM MgCl_2_ at OD_600_ = 0.002 (2e5 CFU mL^−1^) was carefully inoculated to 5-week-old Arabidopsis leaf with a blunt syringe. Plants were held at growth conditions with a transparent plastic dome. Leaves are harvested at 2 days post inoculation (dpi). Twelve independent leaf disks with 4 mm diameter were collected and pooled as one sample (biological repeat). If not specially claimed, each experiment uses 4 samples.

*P. carotovorum* WWP14 was grown in Luria Bertani (LB) media. Five μL of WWP14 solution in 10 mM MgCl_2_ at OD_600_ = 0.02 was dropped on the center of one side of the main vein of 5-week-old Arabidopsis leaves. Plants were held at growth conditions with a sealed transparent plastic dome. The major/minor axes of the infection plague, assumed as an ellipse, were measured by a vernier caliper at 16-24 hours post inoculation (hpi) to determine the plaque area.

*G. cichoracearum* UCSC1 preservation and inoculation method was previously described (Wu et al. 2024). In brief, UCSC1 was maintained on the Arabidopsis *eds1* mutant. Spores from 15∼20 heavily infected *eds1* rosette leaves were evenly spread on 24.5 x 10 inches, and a 48 μm mesh was used to gently brush the inoculum into the leaf. Plants were kept in an inoculation chamber for disease development. For spore counting, 2-3 maximumly extended rosette leaves at ∼8 dpi were collected. Spores were suspended in 0.02% Silwet L-77 plus 0.02% Tween-20 and counted using a LUNA™ Automated Cell Counter. For imaging of infected leaves, plants at ∼10 dpi were used.

### Root commensal bacterial infection vertical plate assay (VPA)

Seeds were surface sterilized by washing with 70% ethanol + 0.01% Triton-X for 3 minutes and then 10% bleach solution for 12 minutes. The seeds were next washed 4 times with sterile DI water, 1 minute each. Sterilized seeds were kept in 0.5 ml sterile water and placed in the dark at 4 °C for 3 days before aseptically transferring to grow on ½-Murashige and Skoog (MS) plate for 7 days under 20°C, 16/8 photoperiod, 60% relative humidity, and light intensity of 100 mE. Seedlings were then aseptically transferred to large square plates with ¼-MS plate previously spread with 150 μL of individual bacterial cultures (OD_600_ = 0.01). Plates were closed with parafilm and placed vertically in the same growth conditions described above; distance between seedlings is more than 2cm. Positions of the vertical plates were rearranged every 2 days to avoid impacts of light direction. Roots architectures are imaged using a scanner at 10 dpi. To determine root colonization for each bacterial isolate at 14 days, all seedlings from an individual vertical plate were aseptically collected in a sterile 5 ml tube, weighed, and carefully washed three times with sterile PBS buffer before homogenization using sterile glass beads in 1 mL sterile PBS. The resulting homogenate was serially diluted and plated on LB plates before incubating at 28°C for 24 hours. The resulting colonies were enumerated and normalized by tissue weight to determine colonization of each bacterial strain.

### Agrobacteria-based transformation

The method for transient expression in *N. benthamiana* was previously described (Lu et al. 2020). Briefly, *A. tumefaciens* GV3101 strains harboring expression constructs were pre-incubated at room temperature in induction media (10 mM MES [pH 5.6], 10 mM MgCl_2_, 150 mM acetosyringone) for 2 h before hand-infiltration into 5-week-old *N. benthamiana* leaves using a 1-mL needleless syringe. Inoculated plants were kept in growth condition. Transformed leaves at 2 dpi were used in downstream studies.

For Arabidopsis floral-dip, Agrobacteria were inoculated into 10 mL LB media with proper antibiotics overnight. After centrifugation, the bacteria were re-suspended in 2 mL of the floral-dip buffer (1/2-MS, 5% sucrose, and 0.02% Silwet L-77). Each flower to be transformed was dipped up-and-down for 10s. The plants with dipped flowers were shaded in full darkness with saturated humidity for 24 h and restore to growth condition for harvest.

### Arabidopsis protoplast transformation

Arabidopsis protoplast preparation method is integrated from previous publications (Yoo et al. 2007; Wu et al. 2009) with optimization presented as follows. In brief, leaf #8/9/10 from 5-week-old Arabidopsis plants were excised and the lower epidermis of each leaf was removed using Scotch tape. The leaves were digested in the enzyme solution for 1.5h. The obtained protoplast was washed in W5 solution, followed by resuspension in MMG solution at a cell concentration of 5×10^5^ protoplast/mL. Each sample of 200 μL protoplast in MMG was gently mixed with less than 20uL plasmid vectors (quantity described below for each individual experiment) followed by 220 μL of PEG solution and kept in darkness for 5 min. Next, 920 μL of W5 solution was added immediately to stop transformation. The transformed sample was washed once in W5 and re-suspended in 1mL W5 solution. Samples were kept under weak light (∼200 lux) for 12h before downstream experimentation. The composition of the solutions noted above is identical to those in Yoo et al. 2007.

### BiFC and quantification

Protoplast samples were transformed with 3.5 μg *pM1089::ADF* and 7 μg *pH1097::WRKY*. After 12h, images containing both BiFC fluorescence and chlorophyll fluorescence were collected by confocal microscopy, with low magnification (10X) and maximum pinhole (∼600 nm) to include large quantities of entire cells. To obtain the most accurate and comparable results, the laser and sensor settings were optimized to avoid saturation and maximize the color depth, which is fixed for the total experiment. Cut-off thresholds defining system ground noise in BiFC and chlorophyll channel respectively are manually determined by mock sample (i.e., pM1089::ADF4 or *pH1097::WRKY29* only). The pixel brightness of BiFC and chlorophyll channels in sample images were globally subtracted by the thresholds and the minus pixels were re-valued to 0. Finally, pixel brightnesses of both channels were summed; BiFC intensity was defined by BiFC signal per chlorophyll signal (reflecting live cells).

### Promoter reporter assay

Protoplast samples were transformed with 6 μg *pBGWφ::pW29/pBAG7/pW46/pW48* (*eGFP* driven by WRKY induced promoter), together w/o 3 μg *p5GWR::WRKY* (WRKY-mCherry) and/or 7μg *p5GWB::ADF* (ADF4-mTagBFP). Images were captured using the same approach as described above for BiFC, with four channels – mTagBFP, eGFP, mCherry, and chlorophyll. Similar to semi-quantitative BiFC, GFP/chlorophyll (above background threshold) was calculated for the general comparison of ADF-WRKY regulated promoter intensity. Other channels were used to inspect whether considerable differences of ADF and WRKY quantities may influence the system.

### ChIP-qPCR

For chromatin immunoprecipitation (ChIP) qPCR, 600 μL of protoplast in MMG solution (appx. 2 × 10^5^ protoplasts) were transformed with 15 μg *p5GWM::WRKY*, with and without 30μg *p5GWH::ADF*. For flg22 treatment, flg22 was used at a final concentration of 1 μM, and was added to the sample 1h before harvesting. After a 12h incubation period, the samples were crosslinked, purified, sonicated, homogenized for IP, using Pierce™ Classic Magnetic IP/Co-IP Kit (Thermo # 88804) following the manufacturer’s instructions. Sonication was performed using a Branson CPX5800 ultrasonicator for a total of 30 min. After overnight immunoprecipitation (IP), followed by removal of the DNA crosslinking, the pulled-down DNA fraction, as well as input samples, were collected using the Zymoclean Gel DNA Recovery Kit (Zymo Research, #4008). Elution buffer (45 μL; not containing EDTA) was used to dissolve the DNA sample for ChIP-qPCR. For each run, 2 μL of eluted DNA was used as template with 3 technical repeats. Related primers are listed in Supplemental Material 5.

### ChIP-seq

For ChIP sequencing (ChIP-seq), the same method used for ChIP-qPCR was followed, yet each 600 μL of protoplast sample was eluted with only 20 μL of elution buffer. The distribution of fragmented DNA size was inspected using a TapeStation system (Agilent 4200) to ensure that the majority of the DNA was between 200-400bp. The library was prepared using the Takara ThruPLEX DNA-Seq Kit (Takara, #R400674) and inspected using the TapeStation. The sequencing was performed using 75 bp paired end mode on an Illumina NEXTSEQ 500.

The analysis of the sequencing output was evaluated for quality control, and sequencing adapters were removed from the reads. Burrows-Wheeler Aligner (BWA) was used to align the reads; results with size >900bp, or quality score < 20, or non-nuclear fragments were eliminated from further analysis. Three biological repeats were pooled together. The peak-calling of the filtered alignments is conducted by MACS3 (Zhang et al. 2008) with default setting, with band width set to 250 and without sub-peaking. ChIP input is used as the control dataset. Single peaks are filtered by such criteria as: (1) they are aligned to promoter regions, (2) both p-value and q-value > 20, and (3) enrichment fold (*EF*) > 2. We define *Total Enrichment Fold* (*TEF*) to evaluate the level of ChIP enrichment on promoters since a promoter may have multiple peaks, as *TEF* = 2^√∑(*log*_2_*EF*)2^. By default, promoters with *TEF* > 3 for A4H group (see Fig. 5A), and *TEF* > 6 for others were defined as enriched promoters, unless otherwise specified. For differential peak analysis and cis-element enrichment analysis, an independent peak-calling was conducted using the same setting, except for enabling sub-peaking to increase resolution. Differential peak analysis was conducted using the MACS3 bdgdiff function. Cis-element enrichment analysis was conducted by MEME-ChIP (Bailey et al. 2015) using the ±250bp of the summit base of each peak, and resulted consensus motifs was profiled using footprintDB (Sebastian and Contreras-Moreira 2014).

### mRNA-seq

Plant RNA was extracted using the RNeasy Plant Mini kit (Qiagen # 74904) following the manufacturer’s instructions. Libraries were prepared using the Roche KAPA HyperPrep RNA Library Preparation Kit with KAPA Unique Dual Index (UDI) adapters following manufacturer’s recommendations. Synthesized libraries passing standard quality control were quantified using a combination of Qubit dsDNA HS and Agilent 4200 TapeStation HS DNA1000 assays. The libraries were pooled in equimolar amounts and quantified using the Invitrogen Collibri Quantification qPCR kit. The pool was loaded onto a single lane of a NovaSeq S4 flow cell, and sequencing was performed in a 2×150bp paired-end format using a NovaSeq 6000 v1.5 300 cycle reagent kit. Base calling was performed using Illumina Real Time Analysis (RTA) v3.4.4. Output of RTA was demultiplexed and converted to FastQ format with Illumina Bcl2fastq v2.20.0.

Data pre-processing, including quality control, adapter trimming, quality filtering, is conducted using FASTP (Chen et al. 2018), with default setting. Reads are counted using Salmon (Patro et al. 2017), with default setting plus --allowDovetail and --recoverOrphans. DEGs are called using DEseq2 (Love et al. 2014), and GO enrichment analysis is conducted through clusterProfiler (Wu et al. 2021). DEseq2 and clusterProfiler are performed in R environments, and others in Python. Additional descriptions are provided in figure legends.

### EPRI algorithm

We innovated a new matrix, *Euclidean Pathway Regulation Index (EPRI)* as a measurement of the regulatory intensity of a biological pathway reflected by DEGs identified from mRNA-seq. The rationale is that the Log2-fold-of-change of all DEGs in a GO pathway (*FC_DEG_*) can create a multi-dimensional linear space, and a sample’s distance to the origin *(Euclidean Pathway Regulation Distance, EPRD)* reflects how different this sample condition is from non-regulated status, assuming each gene in the pathway has equal impact to regulate the strength of the pathway. First, the *EPRD* is defined as:

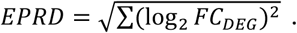

It is obvious that *EPRD* is sensitive to the gene number involved in a pathway. To standardize/normalize the *EPRD*, making this matrix comparable between different biological pathways, we define 1 unit of pathway regulation intensity as the condition when all genes in the pathway are increase/decrease by 2-fold, and make the ratio of *EPRD* to this unit to calculate *EPRI*:

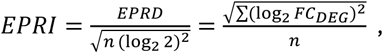

where *n* the total number of genes in the pathway.

### Transcriptional profile analysis on ACTs, ADFs, and WRKYs

Arabidopsis mRNA-seq dataset GSE85932 and GSE151885 were downloaded and decompressed. Raw data was trimmed by Trimmomatic using the default settings to eliminate barcode and low-quality reads. Reads were quantified by Salmon using the same setting as described in mRNA-seq method above. For GSE151885 dataset only, the absolute quantification of all groups was deduced from the 0h group absolute quantification and the relative quantification compared to the 0h group.

### Transcriptional profile analysis on diverse pathogen infection

The mRNA-seq data describing infection process by various pathogens are obtained and analyzed as previously described (Li and Xiao, 2025. Accepted.), with minor modification. Raw counting data of *P. syringae* (basal and ETI), *B. cinerea*, *Hyaloperonospora arabidopsidis*, *Sclerotinia sclerotiorum*, and *Fusarium graminearum* are directly downloaded from Plant Public RNA-seq Database (Yu et al. 2022). The libraries were manually selected and labeled for analysis (Supplemental Material 6). *G. cichoracearum* infection counting was obtained from Li et al., unpublished. Analysis process are described in the corresponding figure legends.

### WRKY immune genotypes references

The information resources of Fig. 3A are listed as below. WRKY4: (Lai et al. 2008). WRKY33: (Birkenbihl et al. 2012). WRKY8: (Chen et al. 2010, 2013; Gao et al. 2013; Ren et al. 2024). WRKY28: (Lin-tao Wu 2011; Gao et al. 2013). WRKY48: (Xing et al. 2008; Gao et al. 2013). WRKY18: (Xu et al. 2006; Pandey et al. 2010). WRKY40: (Pandey et al. 2010). WRKY11: (Journot-Catalino et al. 2006; Jiang et al. 2016). WRKY17: (Journot-Catalino et al. 2006). WRKY22: (Hsu et al. 2013; Kloth et al. 2016). WRKY30: (Zou et al. 2019). WRKY46: (Hu et al. 2012). WRKY70: (Li et al. 2006, 2017; Knoth et al. 2007; Hu et al. 2012; Jiang et al. 2016). WRKY38: (Kim et al. 2008).

## Data availability

TurboID counting (Supplemental Material 1), ChIP-seq enrichment quantification (Supplemental Material 2), mRNA-seq counting (Supplemental Material 3), vector construction process (Supplemental Material4), primer list (Supplemental Material 5), and pathogen infection mRNA library labeling datasets (Supplemental Material 6) will be available upon formal acceptance by a peer-reviewed article. Raw data (readings) of ChIP-seqs and mRNA-seqs will be available at NCBI-SRA at the same time.

## Acknowledgements

We would like to thank Dr. Li Zhang (Soochow University, previously Michigan State University, MSU) for providing technical guidance over CRISPR knocking-out approaches. We would like to thank Dr. Noriko Inada (Osaka Metropolitan University) for generously sharing corresponding plant materials. We would like to thank Dr. Steven Chou (UConn Health, University of Connecticut) for tremendous, unspoken intellectual input related to protein-protein interaction methods. We would like to thank Dr. Sarah Lebeis (MSU) for advice and assistance related to root commensal bacterial infection experiments. We would like to thank Dr. Christopher Contag (MSU) and Dr. Michael H. Bachmann (MSU) for their unspoken material and intellectual support for extensive studies in mammalian expression systems. We would like to thank Dr. Bruce L. Goode (Brandeis University) for the significant intellectual input related to *Hs*DAH. We would like to thank Dr. Huan Chen for technical suggestions on bioinformatics. We would like to thank Jiahui Gong (MSU) for technical guidance over *P. carotovorum* infection assay.

Research at MSU was supported by grants from the National Science Foundation (MCB-1953014) and the National Institutes of General Medical Sciences (1R01GM125743). Research at UMD-IBBR was supported by National Science Foundation (IOS-2224203).

## Reference

1. Bailey TL, Johnson J, Grant CE, and Noble WS. The MEME Suite. Nucleic Acids Research. 2015:43(W1):W39–W49. 10.1093/nar/gkv416

2. Birkenbihl RP, Diezel C, and Somssich IE. Arabidopsis WRKY33 Is a Key Transcriptional Regulator of Hormonal and Metabolic Responses toward Botrytis cinerea Infection. Plant Physiology. 2012:159(1):266–285.

3. Birkenbihl RP, Kracher B, Roccaro M, and Somssich IE. Induced Genome-Wide Binding of Three Arabidopsis WRKY Transcription Factors during Early MAMP-Triggered Immunity. The Plant Cell. 2017:29(1):20–38. 10.1105/tpc.16.00681

4. Branon TC, Bosch JA, Sanchez AD, Udeshi ND, Svinkina T, Carr SA, Feldman JL, Perrimon N, and Ting AY. Efficient proximity labeling in living cells and organisms with TurboID. Nat Biotechnol. 2018:36(9):880–887. 10.1038/nbt.4201

5. Chen H and Day B. Insights into Immune Gene Prediction and Function Through the Evolutionary History of ADF Gene Family. 2024:2024.05.31.596878. 10.1101/2024.05.31.596878

6. Chen L, Zhang L, Li D, Wang F, and Yu D. WRKY8 transcription factor functions in the TMV-cg defense response by mediating both abscisic acid and ethylene signaling in Arabidopsis. Proceedings of the National Academy of Sciences. 2013:110(21):E1963–E1971. 10.1073/pnas.1221347110

7. Chen L, Zhang L, and Yu D. Wounding-Induced WRKY8 Is Involved in Basal Defense in Arabidopsis. MPMI. 2010:23(5):558–565. 10.1094/MPMI-23-5-0558

8. Chen S, Zhou Y, Chen Y, and Gu J. fastp: an ultra-fast all-in-one FASTQ preprocessor. Bioinformatics. 2018:34(17):i884–i890. 10.1093/bioinformatics/bty560

9. Dong C-H and Hong Y. Arabidopsis CDPK6 phosphorylates ADF1 at N-terminal serine 6 predominantly. Plant Cell Rep. 2013:32(11):1715–1728. 10.1007/s00299-013-1482-6

10. Dopie J, Skarp K-P, Kaisa Rajakylä E, Tanhuanpää K, and Vartiainen MK. Active maintenance of nuclear actin by importin 9 supports transcription. Proceedings of the National Academy of Sciences. 2012:109(9):E544– E552. 10.1073/pnas.1118880109

11. Du J, Wang X, Dong C-H, Yang JM, and Yao XJ. Computational Study of the Binding Mechanism of Actin-Depolymerizing Factor 1 with Actin in Arabidopsis thaliana. PLOS ONE. 2016:11(7):e0159053. 10.1371/journal.pone.0159053

12. Gao X, Chen X, Lin W, Chen S, Lu D, Niu Y, Li L, Cheng C, McCormack M, Sheen J, et al. Bifurcation of Arabidopsis NLR Immune Signaling via Ca2+-Dependent Protein Kinases. PLOS Pathogens. 2013:9(1):e1003127. 10.1371/journal.ppat.1003127

13. García-González J and van Gelderen K. Bundling up the Role of the Actin Cytoskeleton in Primary Root Growth. Front Plant Sci. 2021:12. 10.3389/fpls.2021.777119

14. Hsu F-C, Chou M-Y, Chou S-J, Li Y-R, Peng H-P, and Shih M-C. Submergence Confers Immunity Mediated by the WRKY22 Transcription Factor in Arabidopsis. Plant Cell. 2013:25(7):2699–2713. 10.1105/tpc.113.114447

15. Hu Y, Dong Q, and Yu D. Arabidopsis WRKY46 coordinates with WRKY70 and WRKY53 in basal resistance against pathogen Pseudomonas syringae. Plant Science. 2012:185–186:288–297. 10.1016/j.plantsci.2011.12.003

16. Inada N, Higaki T, and Hasezawa S. Nuclear Function of Subclass I Actin-Depolymerizing Factor Contributes to Susceptibility in Arabidopsis to an Adapted Powdery Mildew Fungus. Plant Physiology. 2016:170(3):1420– 1434. 10.1104/pp.15.01265

17. Jaswandkar SV, Katti KS, and Katti DR. Molecular and structural basis of actin filament severing by ADF/cofilin. Computational and Structural Biotechnology Journal. 2022:20:4157–4171. 10.1016/j.csbj.2022.07.054

18. Javed T and Gao S-J. WRKY transcription factors in plant defense. Trends in Genetics. 2023:39(10):787–801. 10.1016/j.tig.2023.07.001

19. Jiang C-H, Huang Z-Y, Xie P, Gu C, Li K, Wang D-C, Yu Y-Y, Fan Z-H, Wang C-J, Wang Y-P, et al. Transcription factors WRKY70 and WRKY11 served as regulators in rhizobacterium Bacillus cereus AR156-induced systemic resistance to Pseudomonas syringae pv. tomato DC3000 in Arabidopsis. J Exp Bot. 2016:67(1):157–174. 10.1093/jxb/erv445

20. Jones JDG, Staskawicz BJ, and Dangl JL. The plant immune system: From discovery to deployment. Cell. 2024:187(9):2095–2116. 10.1016/j.cell.2024.03.045

21. Journot-Catalino N, Somssich IE, Roby D, and Kroj T. The Transcription Factors WRKY11 and WRKY17 Act as Negative Regulators of Basal Resistance in Arabidopsis thaliana. Plant Cell. 2006:18(11):3289–3302. 10.1105/tpc.106.044149

22. Kim K-C, Lai Z, Fan B, and Chen Z. Arabidopsis WRKY38 and WRKY62 Transcription Factors Interact with Histone Deacetylase 19 in Basal Defense. The Plant Cell. 2008:20(9):2357–2371. 10.1105/tpc.107.055566

23. Kloc M, Chanana P, Vaughn N, Uosef A, Kubiak JZ, and Ghobrial RM. New Insights into Cellular Functions of Nuclear Actin. Biology. 2021:10(4):304. 10.3390/biology10040304

24. Kloth KJ, Wiegers GL, Busscher-Lange J, van Haarst JC, Kruijer W, Bouwmeester HJ, Dicke M, and Jongsma MA. AtWRKY22 promotes susceptibility to aphids and modulates salicylic acid and jasmonic acid signalling. J Exp Bot. 2016:67(11):3383–3396. 10.1093/jxb/erw159

25. Knoth C, Ringler J, Dangl JL, and Eulgem T. Arabidopsis WRKY70 Is Required for Full RPP4-Mediated Disease Resistance and Basal Defense Against Hyaloperonospora parasitica. MPMI. 2007:20(2):120–128. 10.1094/MPMI-20-2-0120

26. Kyheröinen S and Vartiainen MK. Nuclear actin dynamics in gene expression and genome organization. Seminars in Cell & Developmental Biology. 2020:102:105–112. 10.1016/j.semcdb.2019.10.012

27. Lai Z, Vinod K, Zheng Z, Fan B, and Chen Z. Roles of ArabidopsisWRKY3 and WRKY4 Transcription Factors in Plant Responses to Pathogens. BMC Plant Biol. 2008:8(1):68. 10.1186/1471-2229-8-68

28. Lappalainen P, Kotila T, Jégou A, and Romet-Lemonne G. Biochemical and mechanical regulation of actin dynamics. Nat Rev Mol Cell Biol. 2022:23(12):836–852. 10.1038/s41580-022-00508-4

29. Li J, Brader G, Kariola T, and Tapio Palva E. WRKY70 modulates the selection of signaling pathways in plant defense. The Plant Journal. 2006:46(3):477–491. 10.1111/j.1365-313X.2006.02712.x

30. Li J, Zhong R, and Palva ET. WRKY70 and its homolog WRKY54 negatively modulate the cell wall-associated defenses to necrotrophic pathogens in Arabidopsis. PLOS ONE. 2017:12(8):e0183731. 10.1371/journal.pone.0183731

31. Li P and Day B. Battlefield Cytoskeleton: Turning the Tide on Plant Immunity. MPMI. 2019:32(1):25–34. 10.1094/MPMI-07-18-0195-FI

32. Li P, Zhang Z, Tong Y, Foda BM, and Day B. ILEE: Algorithms and toolbox for unguided and accurate quantitative analysis of cytoskeletal images. Journal of Cell Biology. 2022:222(2):e202203024. 10.1083/jcb.202203024

33. Li W, Pang S, Lu Z, and Jin B. Function and Mechanism of WRKY Transcription Factors in Abiotic Stress Responses of Plants. Plants. 2020:9(11):1515. 10.3390/plants9111515

34. Lin-tao Wu. Arabidopsis WRKY28 transcription factor is required for resistance to necrotrophic pathogen, Botrytis cinerea. Afr J Microbiol Res. 2011:5(30). 10.5897/AJMR11.781

35. Love MI, Huber W, and Anders S. Moderated estimation of fold change and dispersion for RNA-seq data with DESeq2. Genome Biol. 2014:15(12):550. 10.1186/s13059-014-0550-8

36. Lu Y, Zhang Y, Lian N, and Li X. Membrane Dynamics Regulated by Cytoskeleton in Plant Immunity. International Journal of Molecular Sciences. 2023:24(7):6059. 10.3390/ijms24076059

37. Lu Y-J, Li P, Shimono M, Corrion A, Higaki T, He SY, and Day B. Arabidopsis calcium-dependent protein kinase 3 regulates actin cytoskeleton organization and immunity. Nat Commun. 2020:11(1):6234. 10.1038/s41467-020-20007-4

38. Mair A, Xu S-L, Branon TC, Ting AY, and Bergmann DC. Proximity labeling of protein complexes and cell-type-specific organellar proteomes in Arabidopsis enabled by TurboID. eLife. 2019:8:e47864. 10.7554/eLife.47864

39. Mao Y, Zhang H, Xu N, Zhang B, Gou F, and Zhu J-K. Application of the CRISPR–Cas System for Efficient Genome Engineering in Plants. Molecular Plant. 2013:6(6):2008–2011. 10.1093/mp/sst121

40. Matsumoto T, Higaki T, Takatsuka H, Kutsuna N, Ogata Y, Hasezawa S, Umeda M, and Inada N. Arabidopsis thaliana Subclass I ACTIN DEPOLYMERIZING FACTORs Regulate Nuclear Organization and Gene Expression. Plant and Cell Physiology. 2023:64(10):1231–1242. 10.1093/pcp/pcad092

41. Matsumoto T, Kobayashi K, and Inada N. Arabidopsis thaliana ACTIN DEPOLYMERIZING FACTORs play a role in leaf senescence regulation. 2024:2024.05.14.594232. 10.1101/2024.05.14.594232

42. McDowell JM, Huang S, McKinney EC, An Y-Q, and Meagher RB. Structure and Evolution of the Actin Gene Family in Arabidopsis thaliana. Genetics. 1996:142(2):587–602. 10.1093/genetics/142.2.587

43. Pandey SP, Roccaro M, Schön M, Logemann E, and Somssich IE. Transcriptional reprogramming regulated by WRKY18 and WRKY40 facilitates powdery mildew infection of Arabidopsis. The Plant Journal. 2010:64(6):912–923. 10.1111/j.1365-313X.2010.04387.x

44. Patro R, Duggal G, Love MI, Irizarry RA, and Kingsford C. Salmon provides fast and bias-aware quantification of transcript expression. Nat Methods. 2017:14(4):417–419. 10.1038/nmeth.4197

45. Pollard TD and Goldman RD. Overview of the Cytoskeleton from an Evolutionary Perspective. Cold Spring Harb Perspect Biol. 2018:10(7):a030288. 10.1101/cshperspect.a030288

46. Porter K, Shimono M, Tian M, and Day B. Arabidopsis Actin-Depolymerizing Factor-4 Links Pathogen Perception, Defense Activation and Transcription to Cytoskeletal Dynamics. PLOS Pathogens. 2012:8(11):e1003006. 10.1371/journal.ppat.1003006

47. Qin L, Liu L, Tu J, Yang G, Wang S, Quilichini TD, Gao P, Wang H, Peng G, Blancaflor EB, et al. The ARP2/3 complex, acting cooperatively with Class I formins, modulates penetration resistance in Arabidopsis against powdery mildew invasion. The Plant Cell. 2021:33(9):3151–3175. 10.1093/plcell/koab170

48. Ren C-X, Chen S-Y, He Y-H, Xu Y-P, Yang J, and Cai X-Z. Fine-tuning of the dual-role transcription factor WRKY8 via differential phosphorylation for robust broad-spectrum plant immunity. Plant Comm. 2024:5(12). 10.1016/j.xplc.2024.101072

49. Saile SC, Jacob P, Castel B, Jubic LM, Salas-Gonzáles I, Bäcker M, Jones JDG, Dangl JL, and Kasmi FE. Two unequally redundant “helper” immune receptor families mediate Arabidopsis thaliana intracellular “sensor” immune receptor functions. PLOS Biology. 2020:18(9):e3000783. 10.1371/journal.pbio.3000783

50. Sebastian A and Contreras-Moreira B. footprintDB: a database of transcription factors with annotated cis elements and binding interfaces. Bioinformatics. 2014:30(2):258–265. 10.1093/bioinformatics/btt663

51. Sharma A and Chandran D. Host nuclear repositioning and actin polarization towards the site of penetration precedes fungal ingress during compatible pea-powdery mildew interactions. Planta. 2022:256(2):45. 10.1007/s00425-022-03959-3

52. Shehjar F, Almarghalani DA, Mahajan R, Hasan SA-M, and Shah ZA. The Multifaceted Role of Cofilin in Neurodegeneration and Stroke: Insights into Pathogenesis and Targeting as a Therapy. Cells. 2024:13(2):188. 10.3390/cells13020188

53. Sinha J, Singh Y, and Verma PK. Cytoskeleton remodeling: a central player in plant–fungus interactions. Journal of Experimental Botany. 2024:75(11):3269–3286. 10.1093/jxb/erae133

54. Sun Y, Shi M, Wang D, Gong Y, Sha Q, Lv P, Yang J, Chu P, and Guo S. Research progress on the roles of actin-depolymerizing factor in plant stress responses. Front Plant Sci. 2023:14. 10.3389/fpls.2023.1278311

55. Tanaka K, Takeda S, Mitsuoka K, Oda T, Kimura-Sakiyama C, Maéda Y, and Narita A. Structural basis for cofilin binding and actin filament disassembly. Nat Commun. 2018:9(1):1860. 10.1038/s41467-018-04290-w

56. Teliska LH and Rasband MN. Spectrins. Current Biology. 2021:31(10):R504–R506. 10.1016/j.cub.2021.01.040

57. Tian F, Yang D-C, Meng Y-Q, Jin J, and Gao G. PlantRegMap: charting functional regulatory maps in plants. Nucleic Acids Research. 2020:48(D1):D1104–D1113. 10.1093/nar/gkz1020

58. Tian M, Chaudhry F, Ruzicka DR, Meagher RB, Staiger CJ, and Day B. Arabidopsis Actin-Depolymerizing Factor AtADF4 Mediates Defense Signal Transduction Triggered by the Pseudomonas syringae Effector AvrPphB. Plant Physiology. 2009:150(2):815–824. 10.1104/pp.109.137604

59. Van Ngo H and Mostowy S. Role of septins in microbial infection. Journal of Cell Science. 2019:132(9). 10.1242/jcs.226266

60. Wang J, Lian N, Zhang Y, Man Y, Chen L, Yang H, Lin J, and Jing Y. The Cytoskeleton in Plant Immunity: Dynamics, Regulation, and Function. International Journal of Molecular Sciences. 2022:23(24):15553. 10.3390/ijms232415553

61. Wei M, Fan X, Ding M, Li R, Shao S, Hou Y, Meng S, Tang F, Li C, and Sun Y. Nuclear actin regulates inducible transcription by enhancing RNA polymerase II clustering. Science Advances. 2020:6(16):eaay6515. 10.1126/sciadv.aay6515

62. Wu F-H, Shen S-C, Lee L-Y, Lee S-H, Chan M-T, and Lin C-S. Tape-Arabidopsis Sandwich - a simpler Arabidopsis protoplast isolation method. Plant Methods. 2009:5(1):16. 10.1186/1746-4811-5-16

63. Wu T, Hu E, Xu S, Chen M, Guo P, Dai Z, Feng T, Zhou L, Tang W, Zhan L, et al. clusterProfiler 4.0: A universal enrichment tool for interpreting omics data. Innovation. 2021:2(3). 10.1016/j.xinn.2021.100141

64. Wu Y, Sexton WK, Zhang Q, Bloodgood D, Wu Y, Hooks C, Coker F, Vasquez A, Wei C-I, and Xiao S. Leaf abaxial immunity to powdery mildew in Arabidopsis is conferred by multiple defense mechanisms. Journal of Experimental Botany. 2024:75(5):1465–1478. 10.1093/jxb/erad450

65. Xing D-H, Lai Z-B, Zheng Z-Y, Vinod KM, Fan B-F, and Chen Z-X. Stress- and Pathogen-Induced Arabidopsis WRKY48 is a Transcriptional Activator that Represses Plant Basal Defense. Molecular Plant. 2008:1(3):459–470. 10.1093/mp/ssn020

66. Xing J, Wang Y, Peng A, Li J, Niu X, and Zhang K. The role of actin cytoskeleton CFL1 and ADF/cofilin superfamily in inflammatory response. Front Mol Biosci. 2024:11. 10.3389/fmolb.2024.1408287

67. Xu X, Chen C, Fan B, and Chen Z. Physical and Functional Interactions between Pathogen-Induced Arabidopsis WRKY18, WRKY40, and WRKY60 Transcription Factors. The Plant Cell. 2006:18(5):1310–1326.

68. Yao N, Li J, Liu H, Wan J, Liu W, and Zhang M. The Structure of the ZMYND8/Drebrin Complex Suggests a Cytoplasmic Sequestering Mechanism of ZMYND8 by Drebrin. Structure. 2017:25(11):1657–1666.e3. 10.1016/j.str.2017.08.014

69. Yoo S-D, Cho Y-H, and Sheen J. Arabidopsis mesophyll protoplasts: a versatile cell system for transient gene expression analysis. Nat Protoc. 2007:2(7):1565–1572. 10.1038/nprot.2007.199

70. Yu H, Maoliniyazi M, Han X, Yang H, Zhang Z, Guo Y, Tang X, Li H, Cao Q, Wang S, et al. YUCCA3 interacts with ADF4 to regulate *Arabidopsis* hypocotyl elongation by organizing actin arrays. Plant Physiology and Biochemistry. 2025:223:109877. 10.1016/j.plaphy.2025.109877

71. Yu Y, Zhang H, Long Y, Shu Y, and Zhai J. Plant Public RNA-seq Database: a comprehensive online database for expression analysis of 45 000 plant public RNA-Seq libraries. Plant Biotechnology Journal. 2022:20(5):806– 808. 10.1111/pbi.13798

72. Yuan M, Jiang Z, Bi G, Nomura K, Liu M, Wang Y, Cai B, Zhou J-M, He SY, and Xin X-F. Pattern-recognition receptors are required for NLR-mediated plant immunity. Nature. 2021:592(7852):105–109. 10.1038/s41586-021-03316-6

73. Zhang Y, Liu T, Meyer CA, Eeckhoute J, Johnson DS, Bernstein BE, Nusbaum C, Myers RM, Brown M, Li W, et al. Model-based Analysis of ChIP-Seq (MACS). Genome Biology. 2008:9(9):R137. 10.1186/gb-2008-9-9-r137

74. Zhao S, Jiang Y, Zhao Y, Huang S, Yuan M, Zhao Y, and Guo Y. CASEIN KINASE1-LIKE PROTEIN2 Regulates Actin Filament Stability and Stomatal Closure via Phosphorylation of Actin Depolymerizing Factor. The Plant Cell. 2016:28(6):1422–1439. 10.1105/tpc.16.00078

75. Zou L, Yang F, Ma Y, Wu Q, Yi K, and Zhang D. Transcription factor WRKY30 mediates resistance to Cucumber mosaic virus in Arabidopsis. Biochemical and Biophysical Research Communications. 2019:517(1):118–124. 10.1016/j.bbrc.2019.07.030

