## Supplemental Figures 1-14 for "Actin Depolymerization Factor (ADF) Moonlighting: Nuclear Immune Regulation by Interacting with WRKY Transcription Factors and Shaping the Transcriptome"

**Supplemental Fig. 1: Genotyping of new plant materials generated for this study.**

*adf4-c1*

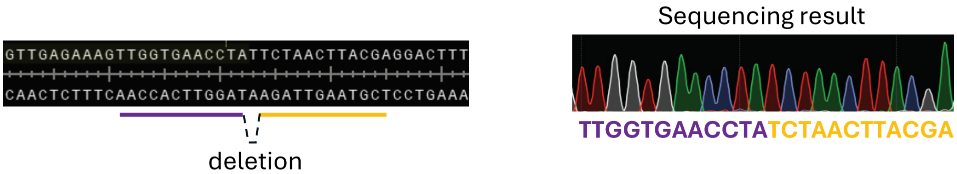


*adf4-2bc*

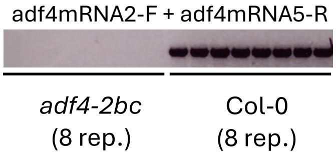


*adf2-c1* (lab internal ID 2g5 #4-3)

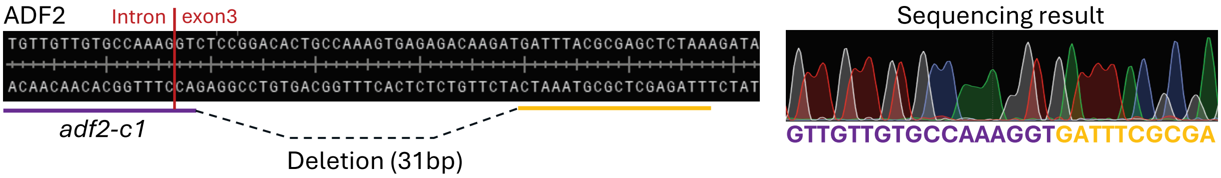


*adf3/adf4* (*adf34*, lab internal ID a34g10 #13-76)

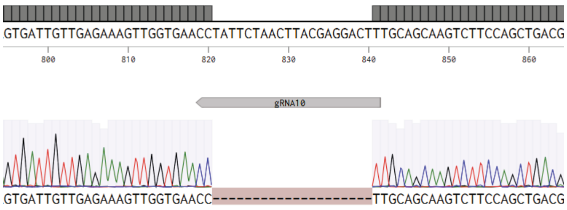


*adf2/3/4* (*a234*), *by adf3/adf4* x *adf2-c1*, F2

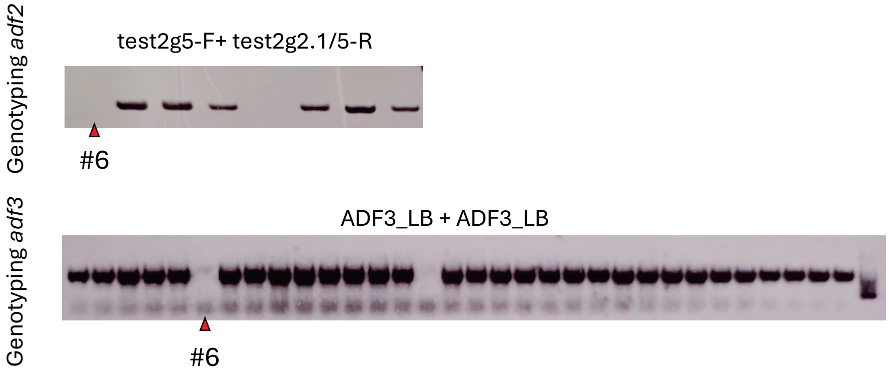


(continuing to next page...)

(continued from previous page)

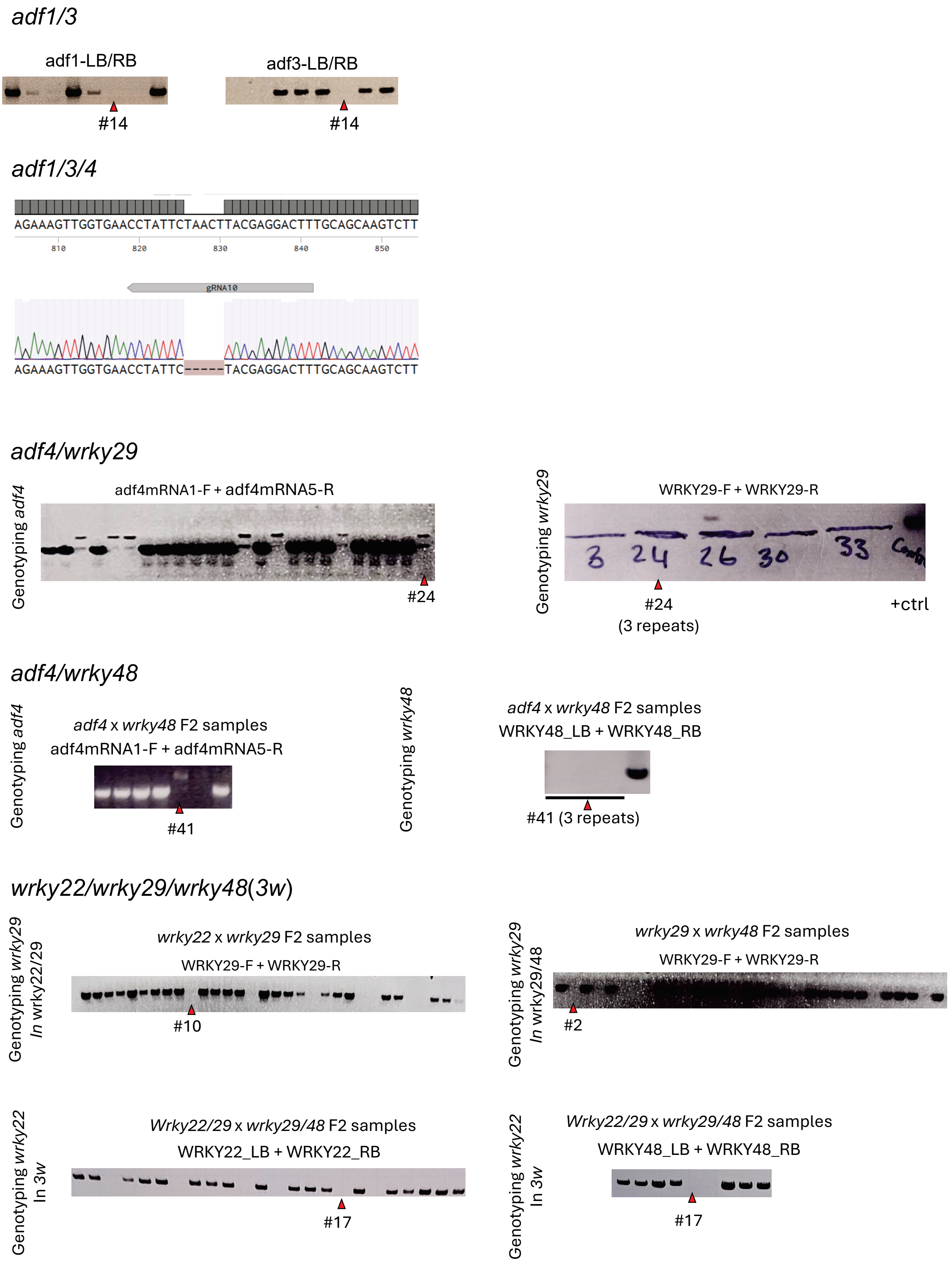


(Figure legends see next page...)

(continued from previous page)

**Supplemental Fig. 1: Genotyping of new plant materials generated for this study.** *adf4-c1* was prepared by CRISPR targeting *ADF4* in Col-0, showing CRISPR T2 line sequencing. *adf4-2bc* was prepared from *adf4-2* by eliminating unexpected insertion on *At1g77500* and then back-crossed to Col-0, showing F2 genotyping PCR. *adf2-c1* was prepared by CRISPR on Col-0, showing CRISPR T2 sequencing. *adf3/adf4* (*a34*) was prepared by CRISPR targeting *ADF4* on *adf3* mutant, showing CRISPR T2 sequencing. *adf2/3/4* (*a234*) was produced by crossing *adf3/adf4* with *adf2-c1*, showing F2 genotyping PCRs. *adf1/3* was prepared by crossing *adf1* with *adf3*, showing F2 genotyping PCR. *adf1/adf3/adf4* (*a134*) was prepared bprepared by crossing *adf4* with *wrky29*, showing F2 genotyping PCR. *adf4/wrky48* was prepared by crossing *adf4* with *wrky48*, showing F2 genotyping PCR. *wrky22/wrky29/wrky48*(*3w*) was prepared by first obtaining two double mutants *wrky22/wrky29* and *wrky29/wrky48* by crossing, which were then crossed again to obtain *3w*, showing all their F2 genotyping y CRISPR targeting *ADF4* in *adf1/adf3* mutant, showing T2 homologous line sequencing. *adf4/wrky29* was results. Red triangle marks lines preserved and used for this study.

**Supplemental Fig. 2: Temporal expression profile of Arabidopsis ACTs upon leaf immune elicitation.**

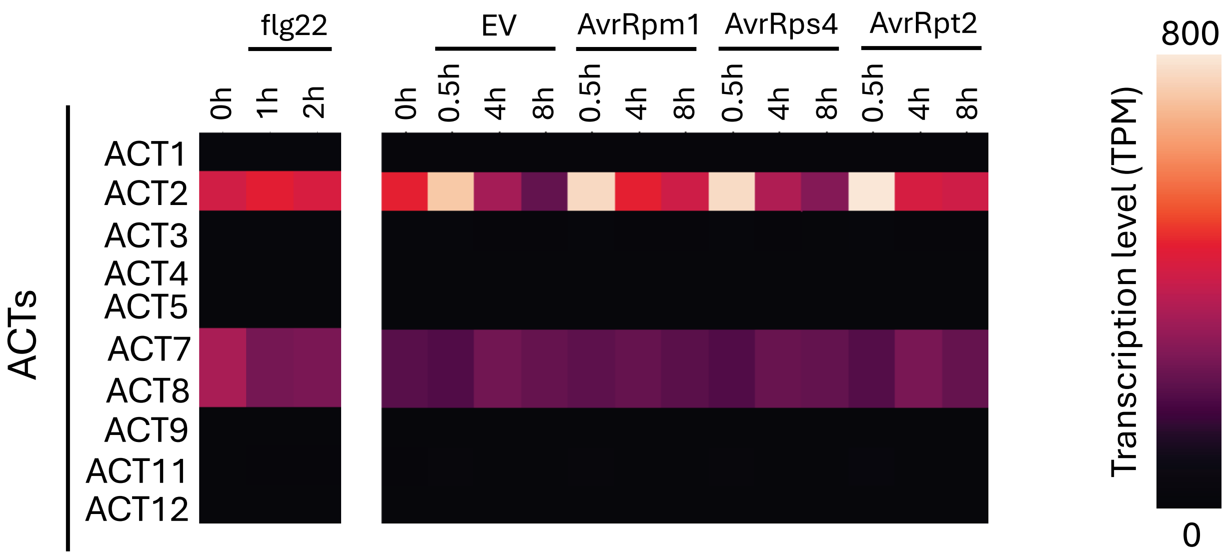


**Supplemental Fig. 2: Temporal expression profile of Arabidopsis ACTs upon leaf immune elicitation.** Same as Fig. 1B, public mRNA-seq datasets GSE85932, describing flg22 treatment and GSE151885, describing DC3000 (w/o avirulent effector) infection were downloaded, processed, and analyzed. Expression levels are represented by TPM. Only ACT2/7/8 are expressed in Arabidopsis leaves. ACT7/8 are stably constitutive; ACT2 are ~2 fold upregulated upon various immune elicitation but fast restored to the resting state. EV, DC3000 with empty vector.

**Supplemental Fig. 3: Temporal expression profile of Arabidopsis WRKYs upon leaf immune elicitation.**

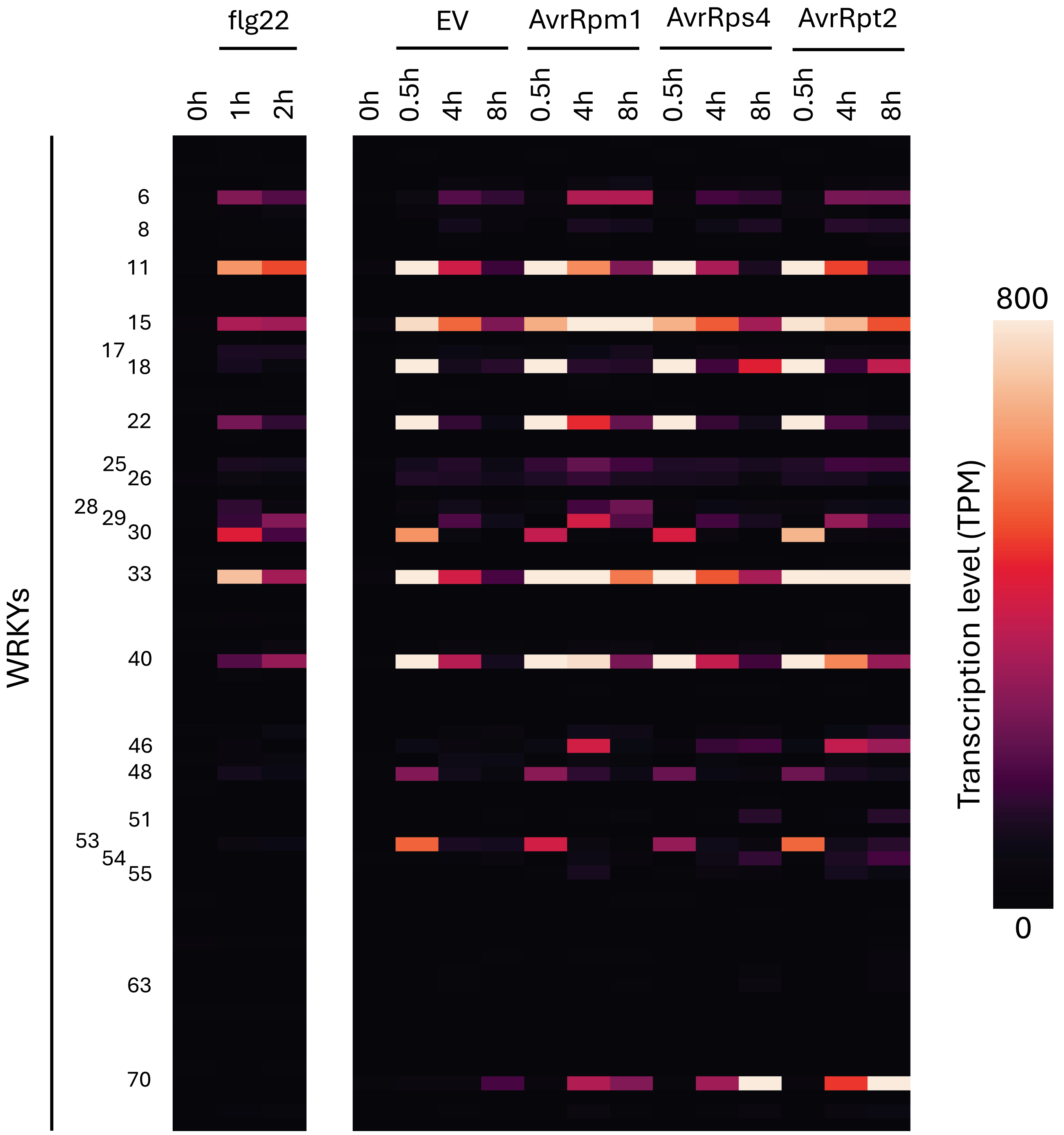


**Supplemental Fig. 3: Temporal expression profile of Arabidopsis WRKYs upon leaf immune elicitation.** Same as Fig. 1B and Supplemental Fig. 2, public mRNA-seq datasets GSE85932, describing flg22 treatment and GSE151885, describing DC3000 (w/o avirulent effector) treatment were downloaded, processed, and analyzed. Expression levels are represented by TPM. WRKYs that are strongly induced by immune elicitation are marked by the number if their ID.

**Supplemental Fig. 4: ADF4$^{d\alpha 3}$ does not interact with actin.**

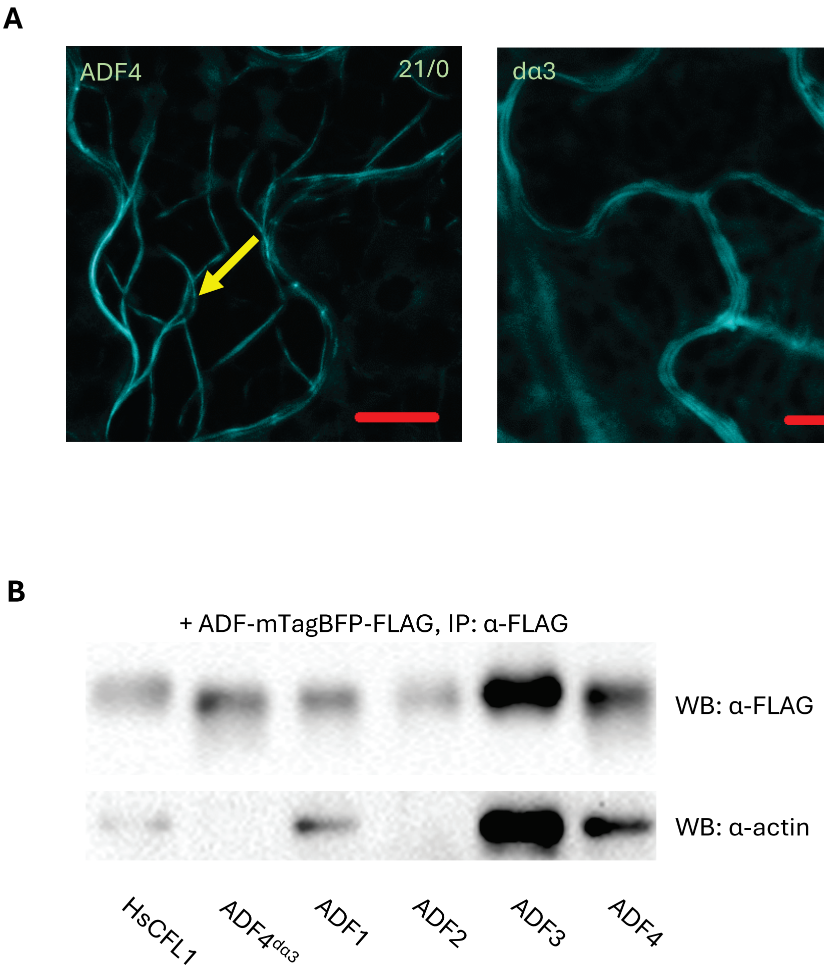


**Supplemental Fig. 4: ADF4$^{d\alpha 3}$ does not interact with actin.** *N. Benthamiana* was infiltrated Agrobacteria containing vector expressing ADF-mTagBFP-FLAG. **A**, subcellular localization of wild type ADF4 and ADF4$^{d\alpha 3}$. ADF4$^{d\alpha 3}$ fails to interact with ACT and therefore cannot attach to actin bundles. Yellow arrow, actin bundle. Numbers of “x/y” on top-right indicates the reproducibility as x: counted cells showing the ADF attaching to actin bundle and y: counts cells that do not. Bar = 10μm. **B**, Co-IP testing the interaction of ADFs to native leaf-expressed NbACTs. ADF4$^{d\alpha 3}$ does not interact with native leaf-expressed NbACTs.

**Supplemental Fig. 5: BiFC assay on WRKY 29 and ADF4 phosphor-mimic mutations.**

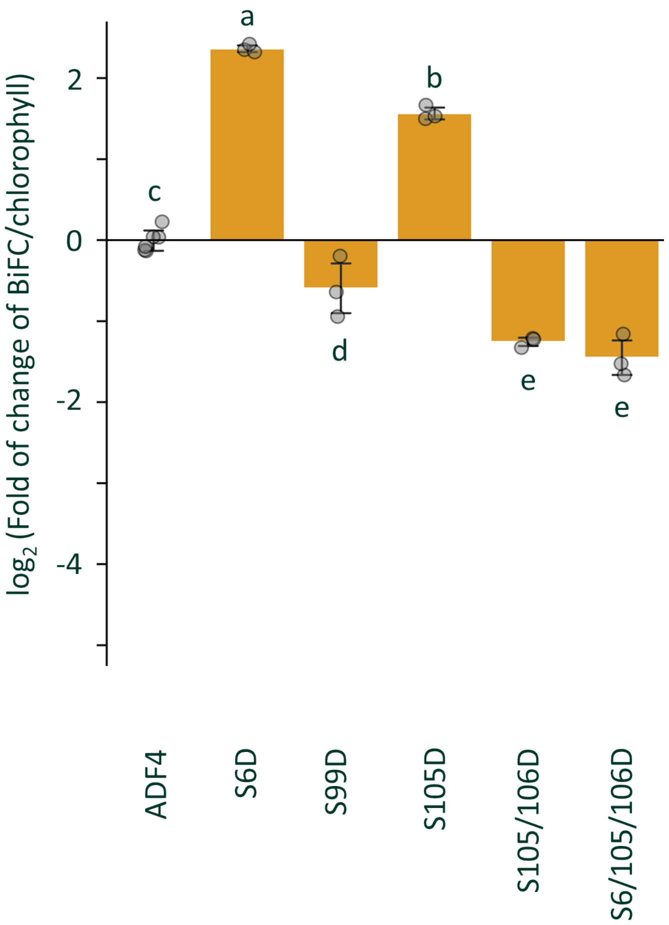


**Supplemental Fig. 5: BiFC assay on WRKY 29 and ADF4 phosphor-mimic mutations.** cY-WRKY29 and equal amounts of nY-ADF4 mutants were transformed Col-0 protoplast for BiFC measurement. The BiFC fluorescence is normalized to log2 fold-of-change compared to wild type ADF4. Phosphorylation can either up- or down-regulate the interaction intensity depending on specific sites. Data passes ANOVA with $P<0.05$; non-overlapping alphabets suggest $P<0.05$ upon post-hoc T-tests adjusted by Benjamini–Hochberg procedure for family-wise error.

### Supplemental Fig. 6: Computation of normalized transcription regulatory effect of ADFs.

**A**

*B*: BFP (ADF) signal — *G*: GFP (pW29 reporter) signal

*R*: RFP (WRKY) signal — *Chl*: chlorophyll (live cell number) signal

$[G]_i = G_i/Chl_i$ ----- GFP per chlorophyll

$[G]_{untriggered} = Mean\ ([G]_i \mid i\ in\ \{-WRKY29, -ADF\})$ ----- pW29 transcription level at naive state

$Reg_i = \frac{G_i}{[G]_{untriggered} * Chl_i}$ ----- hypothetic GFP fold to resting state of each group

$RPW_i = (Reg_i - 1)/R_i$ ----- fold of up-regulation by per unit of WRKY of each group

$RPW_{-ADF} = Mean(RPW_i \mid i\ in\{+WRKY, -ADF\})$ ----- $RPW$ at absence of ADF

$RRPW_i = RPW_i/RPW_{-ADF}$ ----- relative RPW compared to $RPW_{-ADF}$ of individual samples

$ARI_i = (EPW_i - 1)/B_i$ ----- “ADF regulatory index”, or fold of RPW upregulation by per unit of ADF

**B**

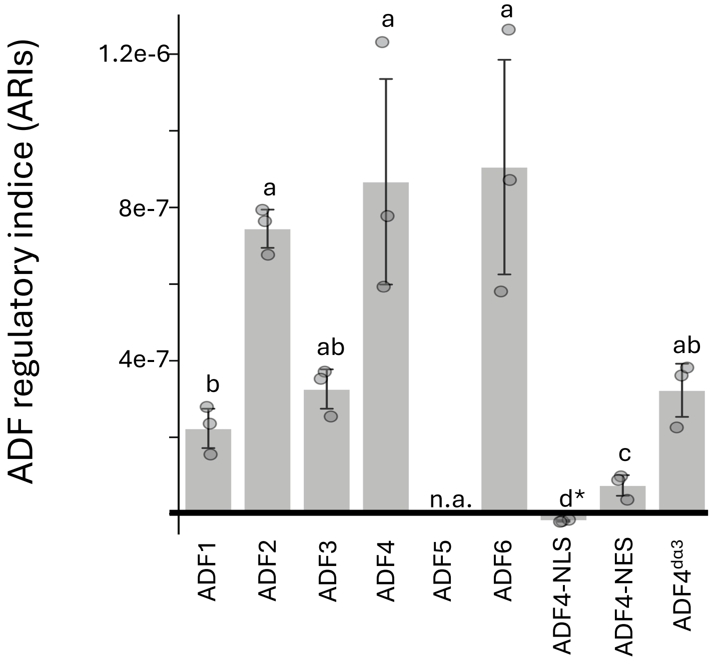


**Supplemental Fig. 6: Computation of normalized transcription regulatory effect of ADFs. A**, description of the calculation algorithm. The normalized transcription regulatory effect of ADFs is defined as ADF regulatory index (ARI). **B**, ARI values of ADFs in Fig. 4F. ADF4-NES, unlike wild type ADF4, shows a low ARI. Data passes ANOVA with $P<0.05$; non-overlapping alphabets suggest $P<0.05$ upon post-hoc T-tests adjusted by Benjamini–Hochberg procedure for family-wise error. n.a., no data generated for ADF5 because reporter signal lower than (–)ADF reference group. *, below 0 because NLS group has highly aggregated, over-saturated signal exclusively in the nucleus, which is not linearly comparable with other groups for this algorithm.

**Supplemental Fig. 8: Bacterial count for the experiment of Fig. 6D-E at 0dpi**

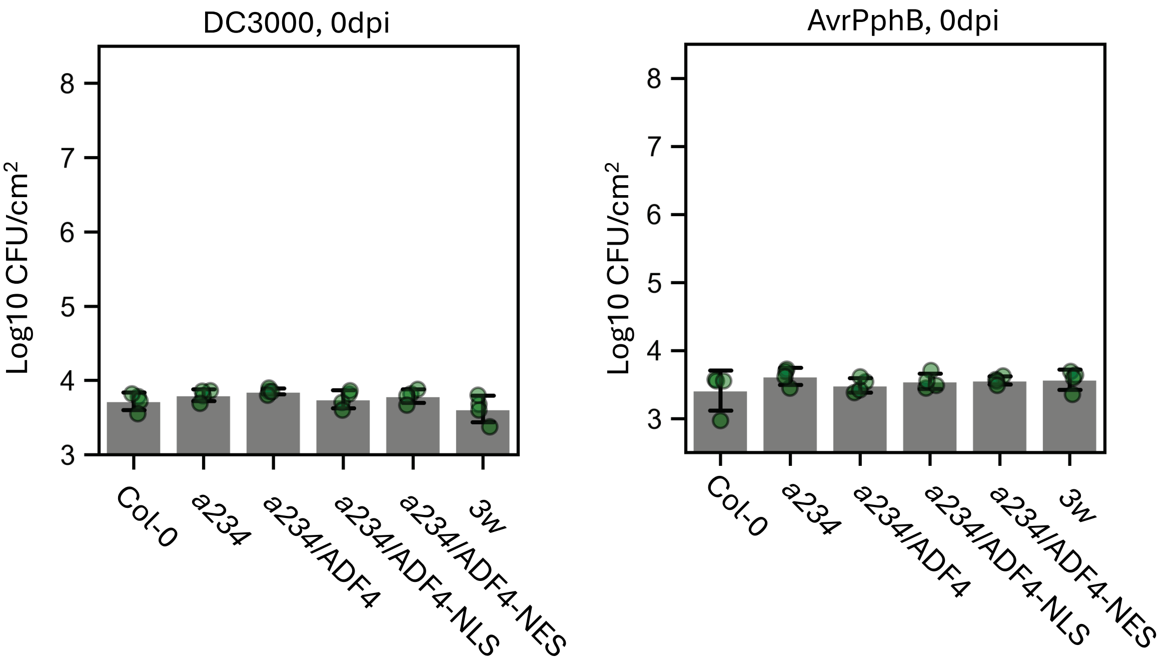


**Supplemental Fig. 8: Bacterial count for the experiment of Fig. 6D-E at 0dpi.** Parallel experiment sets of Fig. 6D-E were immediately sampled and measured after injection. No significant difference is detected.

**Supplemental Fig. 8: DC3000 infection assay using multiple independent ADF4-NES/NLS complementation lines.**

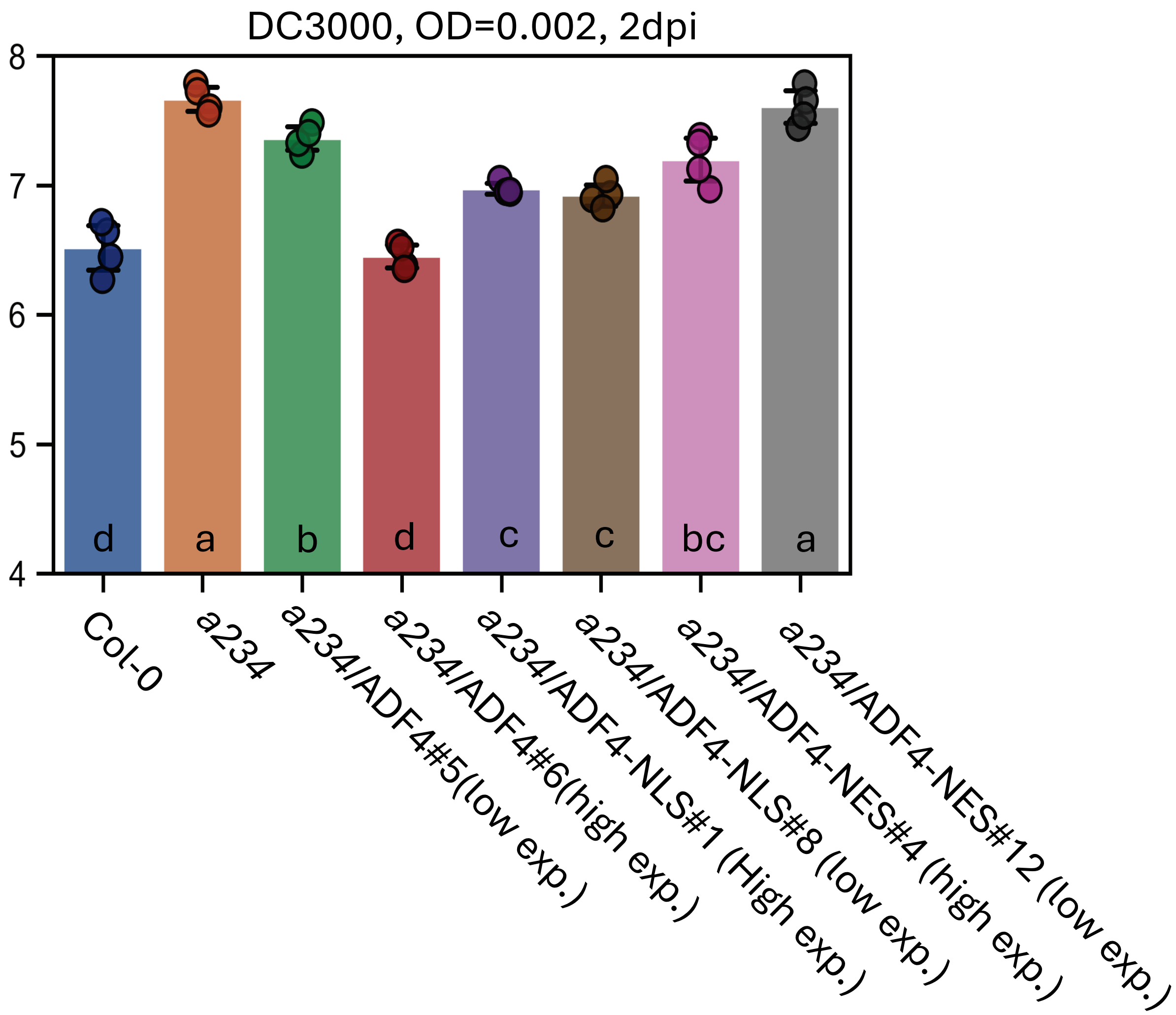


**Supplemental Fig. 8: DC3000 infection assay using multiple independent ADF4-NES/NLS complementation lines.** The experiment setting was the same as Fig. 6D and similar complementation effects of Fig. 6D were observed by multiple, independent lines. Lines with higher ADF4 and ADF4-NES expression have better complementation effect. Data passes ANOVA with $P<0.05$; non-overlapping alphabets suggest $P<0.05$ upon post-hoc T-tests adjusted by Benjamini–Hochberg procedure for family-wise error.

**Supplemental Fig. 9: *a234* and ADF1/2/3/4-silenced line show different phenotype upon** G. cichoracearum **UCSC1 infection.**

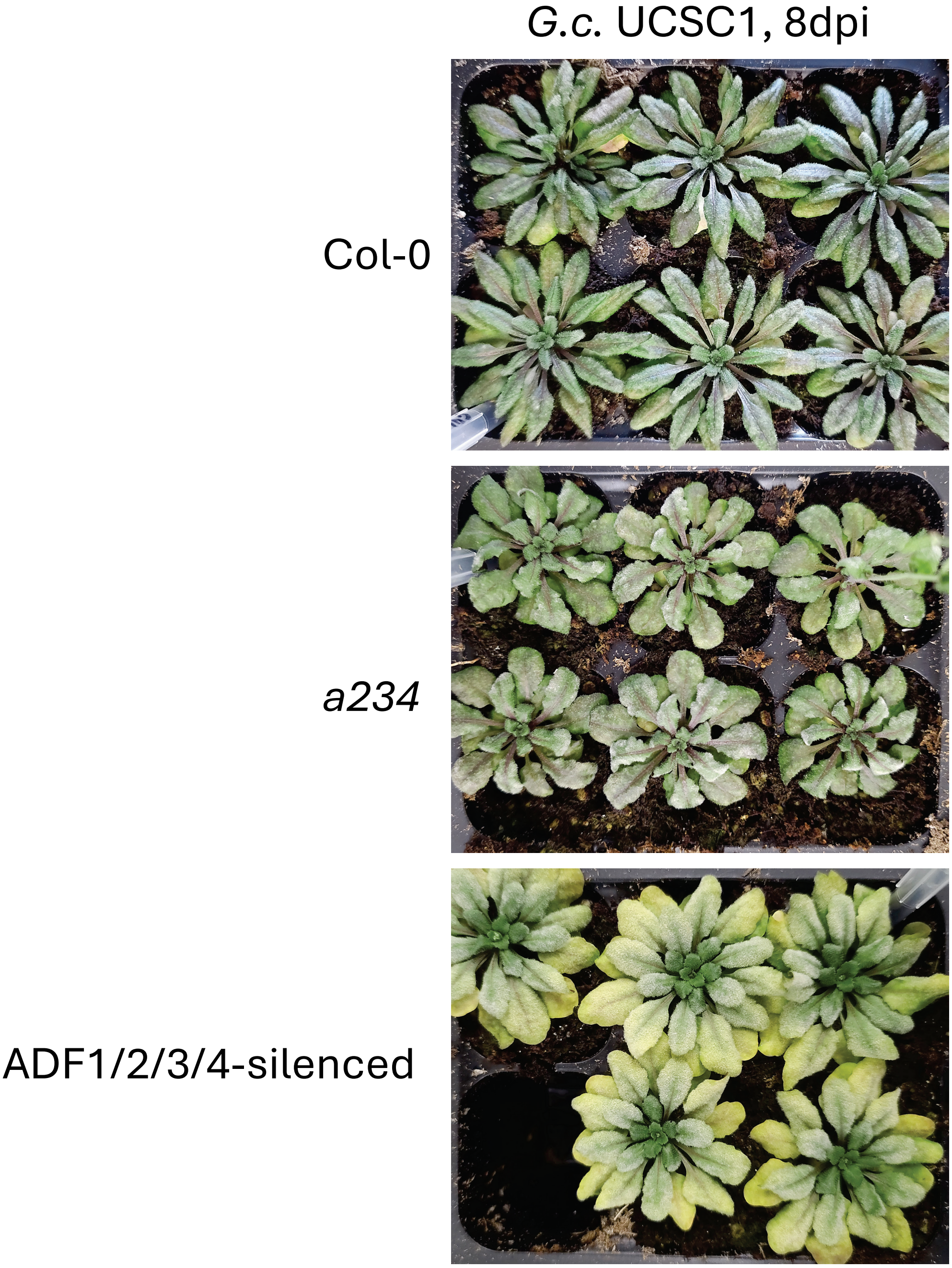


**Supplemental Fig. 9: a234 and ADF1/2/3/4-silenced line show different phenotype upon** G. cichoracearum **UCSC1 infection.** Col-0, *a234*, and ADF1/2/3/4-silenced line were inoculated with *G. cichoracearum* UCSC1 for 8 days. Col-0 and *a234* do not show observable difference at hyphae growth and sporulation level. However, ADF1/2/3/4-silenced lines (Col-0 background) displayed extremely high level of susceptibility accompanied with strong chlorosis.

**Supplemental Fig. 10: The differentiation of gene expression profiles upon infections by different pathogens.**

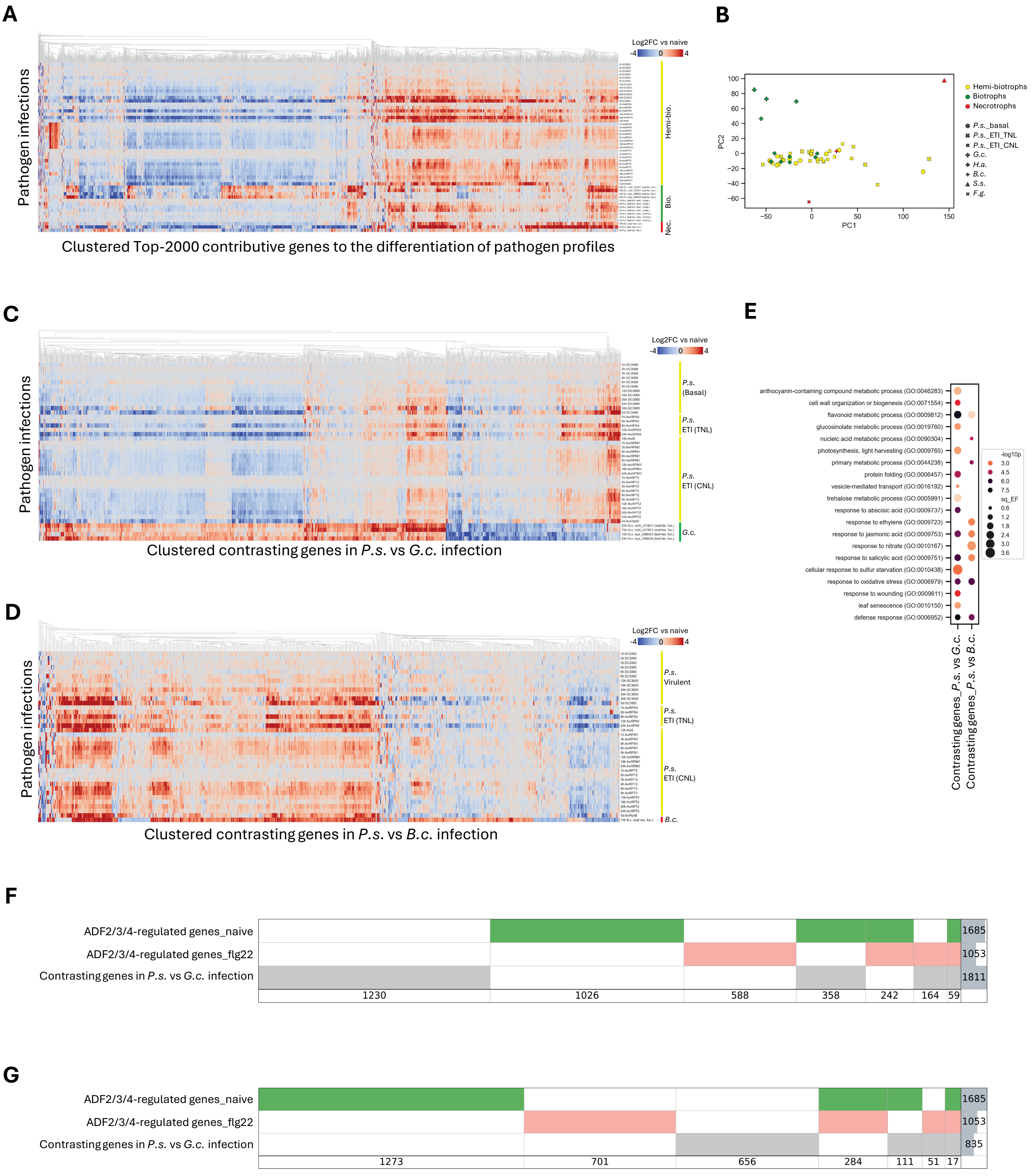


(Figure legends see next page...)

(continued from previous page)

**Supplemental Fig. 10: The differentiation of gene expression profiles upon infections by different pathogens.** mRNA counting data except for those of *G.c.* infection were downloaded from Plant Public RNA-seq Database, manually cleansed and labeled. *G.c.* infection mRNA-seq data were obtained from Li et al., unpublished. FPKM baseline is set to 0.5 to force logarithmic comparison. **A**, Clustered gene expression pattern of different biotrophic, hemi-biotrophic, and necrotrophic pathogen infections. Gene Log2 fold-of-change (Log2FC) compared to corresponding naive/mock datasets is used to perform principal component analysis (PCA). Genes exhibiting top-2000 contribution to the principal components of pattern differentiation are displayed as clustered heatmap. Biotrophs, hemi-biotrophs, and necrotrophs show obviously different clustering patterns. **B**, PCA results (2D) of different pathogen infection datasets. Powdery mildew (*G.c.*) and Botrytis (*B.c.*) show very different gene expression pattern compared to *P.s.* DC3000. **C**, Clustered expression heatmap of genes showing contrast patterns in *P.s.* and *G.c.* infection. “Genes of contrasting pattern” is defined as genes with an absolute difference greater than 1.6 between the mean Log2FC-vs-naïve of all *P.s.* datasets and the mean of datasets of the other pathogen. 1811 contrasting genes are found, which describe a different expression profile between *P.s.* and *G.c.* infection. **D**, Clustered expression heatmap of genes showing contrast patterns between *P.s.* and *B.c.* infection. Several clusters, among the 835 contrasting genes, show different patterns between *P.s.* and *B.c.* infection. **E**, GO pathway enrichment analysis of *P.s.* vs *G.c. and P.s.* vs *B.c.* contrasting genes. The genes showing different trends between different infection processes are enriched in pathways regulating defense compound metabolism, immune response, hormone signaling, etc., and results of *P.s.* vs *G.c.* and *P.s.* vs *B.c.* show different patterns of enrichment. **F**, supervene diagram describing the overlapping level of ADF2/3/4-regulated genes (Fig. 7) and genes showing contrasting pattern between *P.s.* vs *G.c.* infection. Only 31% of the contrasting genes are regulated by ADF2/3/4. **G**, supervene diagram describing the overlapping level of ADF-regulated genes (Fig. 7) and genes showing contrasting pattern between *P.s.* vs *B.c.* infection. Only 21% of the contrasting genes are regulated by ADF2/3/4. **F** and **G** suggest that ADF2/3/4 do not regulate the majority of genes signaturing the difference of *G.c.* and *B.c.* infection compared to that of *P.s.*. Therefore, it is reasonable to observe that *a234* and ADF4 complementation lines show different immune phenotypes on these three pathogens. Abbreviations: *B.c.*, *Botrytis cinerea*; *F.g.*, *Fusarium graminearum*; *G.c.*, *Golovinomyces cichoracearum*; *H.a.*, *Hyaloperonospora arabidopsidis*; *S.s.*, *Sclerotinia sclerotiorum*.

**Supplemental Fig. 11: The correlation matrix of mRNA counting between samples produced by this study.**

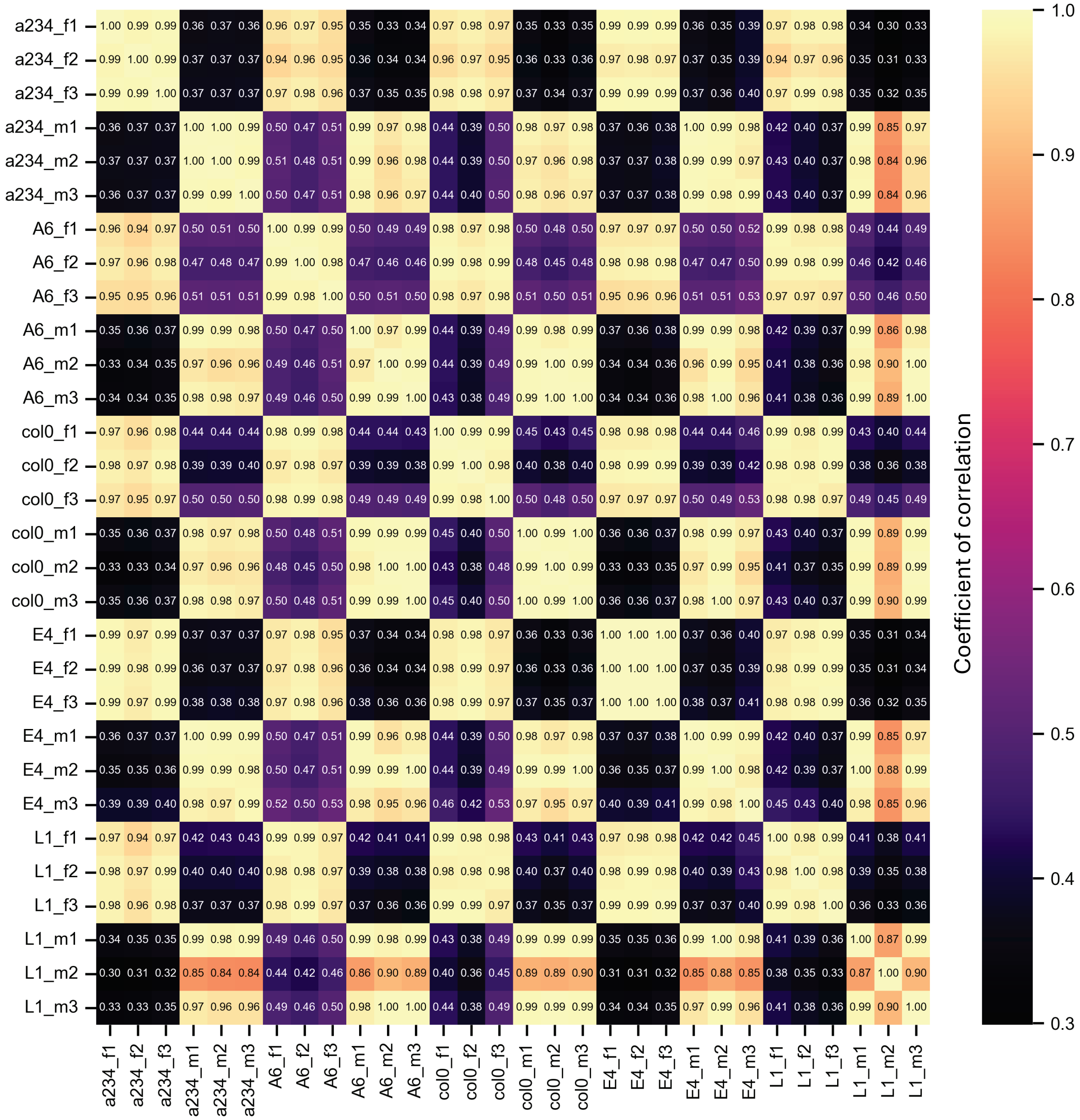


**Supplemental Fig. 11: The correlation matrix of mRNA counting of samples produced by this study.** The correlation efficient of between all datasets of Fig. 7 are calculated. All samples except for sample L1_m2 show satisfying reproducibility. Sample L1_m2 (i.e., *a234*/ADF4-NLS #1-rep2) did not show good reproducibility compared to the other two repeats. After inspection, we found this sample has poor sequencing quality and low matching rate, so we abandoned this sample for further analysis.

**Supplemental Fig. 12: The SuperVenn diagram describing the overlapping levels of PAMP-regulated genes and ADF-regulated genes.**

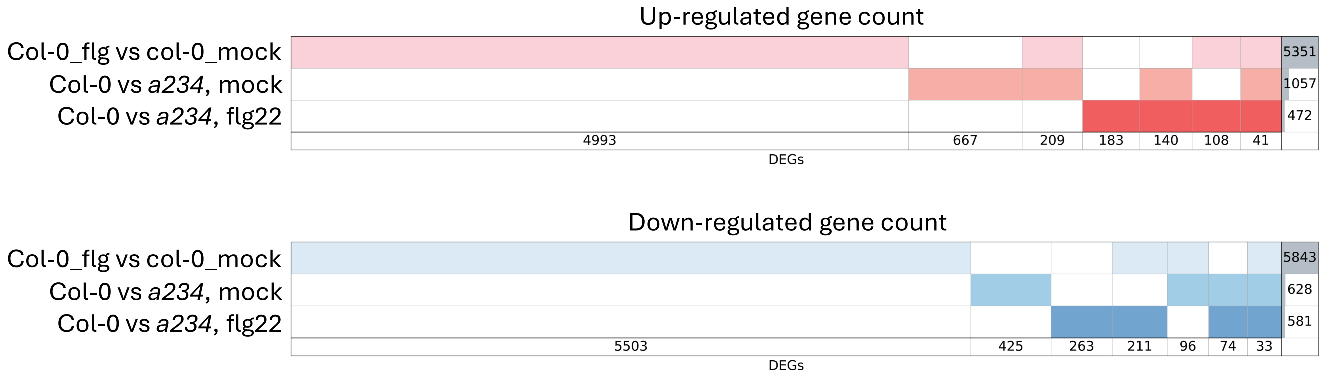


**Supplemental Fig. 12: The SuperVenn diagram describing the overlapping levels of PAMP-regulated genes and ADF-regulated genes.** 498 DEGs up-regulated by ADFs (37% of total) are also up-regulated by flg22; 414 DEGs down-regulated by ADFs (46% of total) are also down-regulated by flg22, both of which indicate that ADFs are heavily involved in PTI and that lack of ADFs may dampen the robustness of immune signaling.

**Supplemental Fig. 13: All-in-one SuperVenn diagram describing the overlapping levels of all DEGs in Fig. 7.**

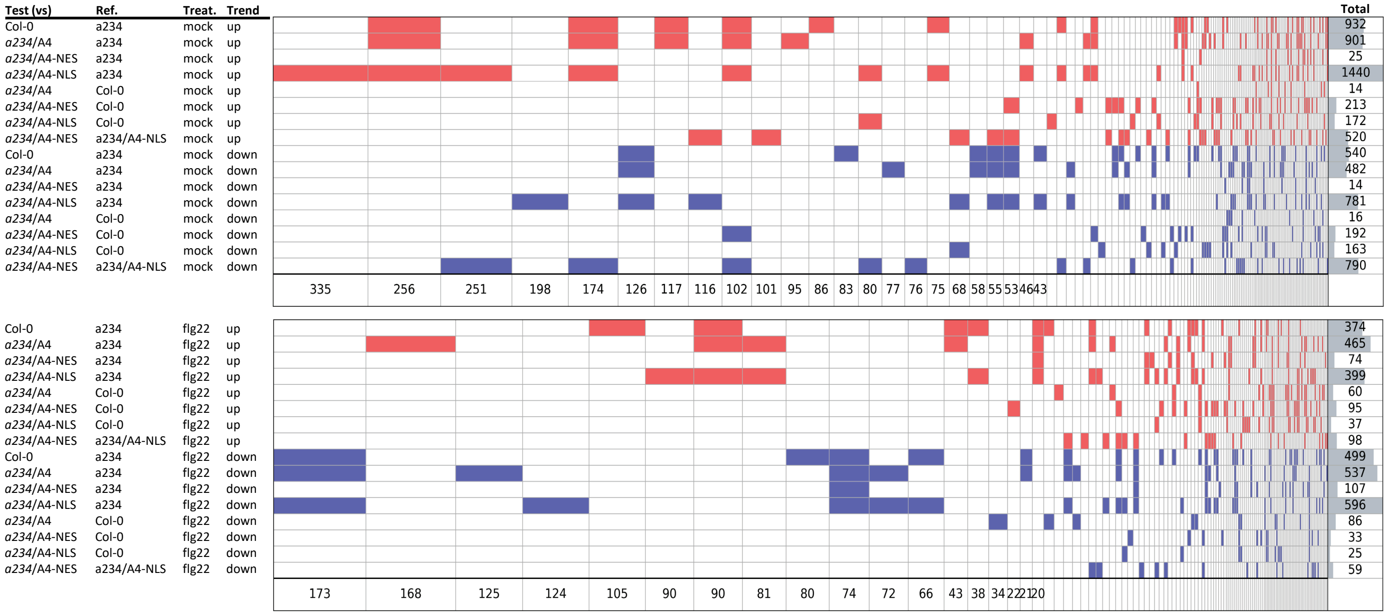


**Supplemental Fig. 13: All-in-one SuperVenn diagram describing the overlapping levels of DEGs in Fig. 7.** This figure serves to provide additional information for interested readers for further interpretation.

**Supplemental Fig. 14: ADF-up/downregulated genes are involved in different signaling pathways.**

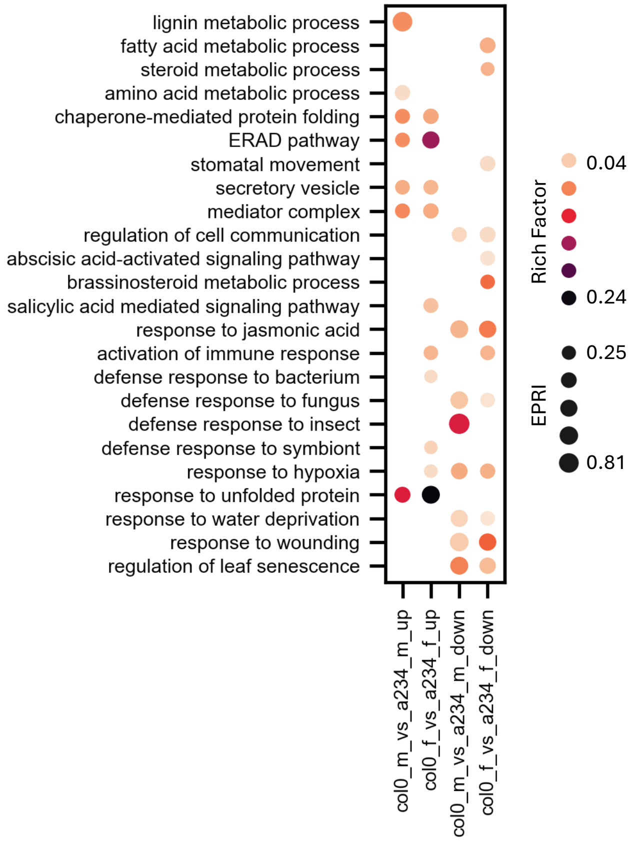


**Supplemental Fig. 14: ADF-up/downregulated genes are involved in different signaling pathways.** The ADF-up/downregulated DEGs, at resting state or elicited, are split for respective GO pathway enrichment analysis. ADF-upregulated genes are enriched in lignin metabolism, protein folding, vesicle secretion, and (transcription-related) mediator complex at the resting state. Upon PAMP elicitation, ADF-upregulated DEGs are additionally enriched in defense response and SA signaling. ADF-downregulated DEGs are generally involved in JA and JA-related defense response, such as response to wounding, insect, and fungus. Upon PAMP elicitation, additional pathways related to ABA and brassinosteroids are emerged.
